# Cell-type-specific ATF6α programs regulate epithelial mitochondrial homeostasis and pericyte remodeling during physiological and exposure-accelerated lung aging

**DOI:** 10.64898/2026.07.09.737329

**Authors:** Xuejian Huang, Jonathan E. Bard, Bat-Ider Tumenbayar, Kokilavani Vedagiri, Caryn E. Nelson, Harshavardhan Kenche, Celia E. Reynolds, Adriana S. Leme, Stephen J. Moore, Noa A. Perry, Steven D. Shapiro, Yaron Perry, Yongho Bae, Anna Blumental-Perry

## Abstract

Proteostasis declines with lung aging, while the role of the Unfolded Protein Response (UPR) in lung aging and age-associated pulmonary diseases remains understudied. We investigated how deficiency in the UPR sensor ATF6α affects physiological and smoke exposure–accelerated lung aging. ATF6α -deficient mice exhibited accelerated alveolar simplification, a sign of lung parenchymal aging, which was exacerbated by smoking. Nevertheless, small airway vascular fibrotic remodeling, a prominent smoking-induced pathology, was not evident in smoke-exposed ATF6α -deficient mice. Mechanistically, these divergent phenotypes arose from cell-type-specific ATF6α programs. In alveolar epithelial type 2 cells (AEC2s), the facultative progenitors of the lung parenchyma, ATF6α maintained mitochondrial bioenergetics and sustained efficient redifferentiation into alveolar epithelial type 1 cells (AEC1s). In lung pericytes, ATF6α promoted extravasation, redifferentiation into myofibroblast-like cells, and production of collagens 1 and 3. These findings identify ATF6α as a cell-type-specific regulator of differentiation programs during lung aging and highlight the need to study ATF6α under defined physiological and pathological contexts before therapeutically targeting this pathway.

## Introduction

The lung contains one of the body’s largest epithelial surfaces and is continuously exposed to inhaled environmental stressors. To preserve function, diverse lung-resident cells rely on robust repair, immune, and stress-response programs (1). With aging, these adaptive mechanisms progressively fail. Lung aging is marked by reduced tissue elasticity, weakening of respiratory muscles, diminished immune function, and impaired repair capacity of lung stem cells (2). This decline begins around age 35 and proceeds at approximately 1% loss of total lung capacity per year (3). Cumulative exposure to toxins, including smoke and pollution, further accelerates this process (4–13).

The most prominent structural feature of physiological lung aging is alveolar simplification, which presents as enlarged alveolar airspaces caused by loss of terminally differentiated **a**lveolar **e**pithelial type **1 c**ells (**AEC1s**) and exhaustion of regenerative **a**lveolar **e**pithelial type **2 c**ells (**AEC2s**), the distal lung facultative progenitors (14). In enlarged alveoli, gas exchange becomes less efficient because increased alveolar diameter creates a mismatch between ventilation and vascular perfusion. This structural and cellular deterioration reduces oxygen delivery to tissues and contributes to age-related decline in pulmonary function.

Inhaled toxins accelerate alveolar simplification, with tobacco use being the largest contributor to accelerated lung aging. Compared with non-smokers, smokers often show a 20-30-year advancement of lung aging and are at increased risk of **C**hronic **O**bstructive **P**ulmonary **D**isease (**COPD**), a disease characterized by cough, shortness of breath, and progressive premature decline in lung function (15). COPD is the most prevalent respiratory disease worldwide, affecting nearly 16 million adults in the U.S. and 213.4 million people globally (16). CS-induced COPD accounts for 70% of COPD cases in high-income countries, disproportionately affects older adults, and remains a major cause of disability, hospitalization, with limited therapeutic options (17). COPD pathogenesis is complex and involves exhaustion of AEC2 repair potential, persistent inflammation, oxidative stress, protease/anti-protease imbalance, and destruction of the lung parenchymal **e**xtra**c**ellular **m**atrix (**ECM**) (18, 19). **S**mall **a**irways and **v**asculature **f**ibrotic **d**isease (**SAVFD**), defined by fibrosis of small airways and their associated blood vessels, represents one of the earliest pathological changes in COPD progression (20–24). SAVFD is believed to disrupt homeostasis at the interface of small airways, lung parenchyma, and associated vasculature, contributing to alveolar loss and accelerated alveolar simplification. Because alveolar simplification is currently considered irreversible, and because no lung rejuvenation approaches or SAVFD-directed therapies are available, defining the drivers of these pathological remodeling processes remains a critical unmet need.

Loss of proteostasis and chronic maladaptive stress responses are hallmarks of aging and contributors to diseases with age-related onset. In aging cells, declining **e**ndoplasmic **r**eticulum (**ER**) folding capacity and variable induction of the **u**nfolded **p**rotein **r**esponse (**UPR**) are well documented (25). However, how ER proteostasis failure and UPR signaling contribute to lung aging and aging-associated remodeling remains understudied (26–28).

UPR coordinates a transcriptional program that enhances ER function, protein folding capacity, quality control, and antioxidant defense. This response is mediated by three conserved sensors: **p**rotein kinase RNA-like **e**ndoplasmic **r**eticulum **k**inase (**PERK**), **a**ctivating **T**ranscription **F**actor **6** (**ATF6**), and **i**nositol-**r**equiring **e**nzyme **1α** (**IRE1α**) (29). We and others have shown that CS and other harmful exposures trigger UPR, primarily through PERK activation (27, 30–32). We also showed that exposure of naïve mice to a single cigarette activates the ATF6α branch, especially in AEC2s (27). Function of ATF6α branch during ER homeostasis maintenance and involvement in various physiological processes, such as aging, that extend beyond the traditional toxin induced UPR are not well defined. Based on these findings, we investigated how the UPR sensor and latent transcription factor ATF6α, referred to here as ATF6, contributes to physiological and cigarette smoke exposure-accelerated lung aging (33, 34).

ATF6 is a type II transmembrane ER glycoprotein that remains inactive at the ER membrane until stress-induced trafficking to the Golgi, where site-1 and site-2 proteases cleave it to release an active transcription factor fragment (35, 36). During acute stress, ATF6 signaling is generally protective because it supports rapid ER folding adaptation and promotes recovery from injury (37–41). However, chronic ATF6 activity may also have detrimental effects, particularly in the lung (42–50). Because lung parenchyma contains approximately 45 distinct cell identities, including AEC1s, AEC2s, endothelial cells, fibroblasts, pericytes, and macrophages, UPR outcomes are unlikely to be uniform across all lung cell types (51). Emerging evidence supports this view, suggesting that UPR signaling is cell-type specific, tailored to the metabolic and physiological demands of each cell population, and depends on repertoire of transcription factors available in specific cell types to interact with ATF6 (33, 34, 52).

Here, we define cell-type-specific mechanisms by which ATF6 fine-tunes lung responses to physiological aging and smoke-induced stress. We show that ATF6 is required to maintain the redifferentiation capacity of AEC2s and lung pericytes, with divergent consequences: protection from premature alveolar simplification in the alveolar epithelium, but promotion of SAVFD in CS-exposed lungs.

## Results

### Absence of ATF6 accelerates alveolar simplification in global ATF6-KO mice

To determine whether ATF6 deficiency affects lung aging, we compared lung phenotypes of 8-and 20-month-old **w**ild-**t**ype (**WT**) and global ATF6-deficient (**ATF6-KO**) mice (53).

We observed an increase in terminal deoxynucleotidyl **t**ransferase d**U**TP **n**ick-**e**nd **l**abeling (**TUNEL**)-positive cells (**Fig. 1A and B**) and a statistically significant increase in the average size of alveoli (evaluated as longer **A**verage **C**hord **L**ength (**ACL**) compared to WT mice), which is indicative of alveolar simplification (**Fig. 1C, D and F**). Age-related alveolar simplification was further supported by comparing ACLs in 8-and 20-month-old mice, with an increase of 3.98 +/- 2.5 μm for WT (p=0.004, **Fig. 1E**) and 5.6 +/-3.8 μm for ATF6-KO mice (p=0.004, **Fig. 1G**). There was a 35% reduction in the number of macrophages/μm of alveolar walls (**Fig. 1H and I**). This anti-inflammatory effect of ATF6 deficiency may be expected because of previously reported proinflammatory effects of ATF6 (54–56). Overall, our findings suggest that ATF6 deficiency accelerates cell loss and moderate airspace enlargement without increased inflammation, a pattern observed in aging human lungs (6, 57, 58). There was no significant difference between WT and ATF6-KO mice in collagen accumulation between small airways and blood vessels that supply them (**Fig. 1J and K**).

**Figure 1.**
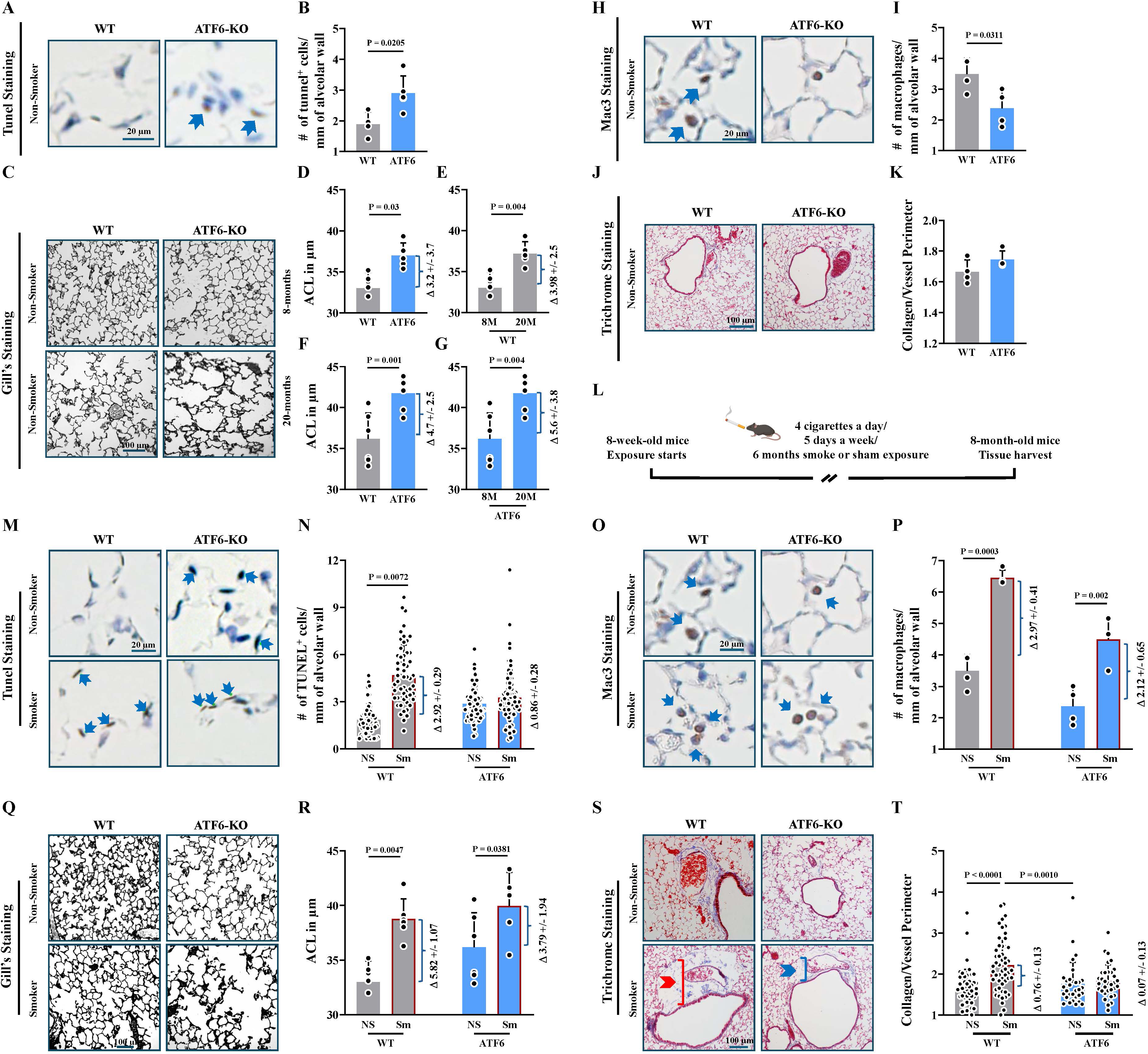
ATF6 deficiency increases apoptosis, accelerates age- and exposure-related alveolar simplification, but prevents SAVFD in mice smokers. Lungs were collected from age-matched WT (grey graphs) and ATF6-KO (blue graphs) mice at 8 months of age, unless indicated otherwise. (**A - B**) TUNEL staining; arrows point to apoptotic nuclei identified by brown staining. (**A**). Graph demonstrates quantification of the number of TUNEL-positive cells per mm² of alveolar area; (10 fields/animal, n=4 for WT; n=6 for ATF6-KO) (**B**). (**C - G**) Gill’s staining. Upper panels show lungs of 8-month-old mice, and lower panels show lungs from 20- month-old mice, respectively. Asterisks mark emphysematous areas. (**C**). Graphs demonstrate quantification of average ACL in 8-month-old (**D**) and 20-month-old (**F**) WT and ATF6-KO mice. (**E**) and (**G**) show age-dependent changes in ACL within the same genotypes (8-12 fields/lung, n=5 for WT and n=6 animals for ATF6-KO). (**H - I**) Macrophage-specific staining by anti-Mac3 immunohistochemistry (**IHC**) (**H**). Graph demonstrates quantification of the number of Mac3^+^ cells per mm² of alveolar area. (10 field/animal, n=3 for WT; n=4 for ATF6- KO) (**I**). (**J-K**) Masson’s Trichrome collagen staining (**J**). Graphs on the left demonstrate quantification of the collagen/vessel perimeter ratios. (calculated from the sectioned airways, usually 25-30 airways/animal, n=4 for WT; n=3 for ATF6-KO) (K). In the quantitative analyses above, each dot represents an individual animal. (**L**) Schematic of the experimental design and CS exposure protocol. WT and ATF6-KO mice were maintained as non-smokers (NS) or exposed to CS as smokers (Sm) starting at 8 weeks of age (four cigarettes per day, five days per week) for six months to model lung changes seen in human smokers and COPD patients. (**M - N**) Apoptosis in lung tissues was evaluated by TUNEL staining. (**M**) Representative images of TUNEL staining; arrows point to brown nuclei positive for fragmented DNA. Graphs demonstrate the number of TUNEL^+^ cells per mm of alveolar wall; each dot represents one field of view, where 10 fields were chosen serendipitously and analyzed for each animal, n=4 for WT NS and Sm and n=5 animals for ATF6-KO NS and Sm (**N**). (**O - P**) Representative images of Mac3 staining; arrows point to brown Mac3^+^ Macrophages. (**O**) Graphs demonstrate the number of Mac3^+^ cells per mm of alveolar wall; each dot represents one field of view, with 10 fields chosen serendipitously and analyzed for each animal, n=4 for all groups (**P**). (**Q - R**) ACL was used as a morphometric measure of airspace enlargement. (**Q**) Representative images of lung parenchyma. (**R**) Quantification of ACL; each dot represents one animal, with n=6 for WT NS, n=7 for WT-Sm, n=6 for ATF6-KO NS, and n=5 for ATF6- KO Sm, 10 fields chosen serendipitously and analyzed for each animal. (**S - T**) Fibrotic changes were visualized by Masson’s Trichrome staining. (**S**) Representative images; red and blue brackets indicate corresponding perivascular and peribronchiolar regions in WT and ATF6-KO Sm, respectively. Please note the significant perivascular and peribronchiolar collagen deposition detectable around terminal bronchioles and their corresponding blood vessels in WT smoker mice but not in ATF6-KO smoker mice. (**T**) Graphs demonstrate quantification of collagen deposition, estimated as the collagen/vessel perimeter ratio. A total of 25-33 small airways and their corresponding blood vessels were scored per animal, with n=4 animals for WT NS, n=6 for WT-Sm, n=3 for ATF6-KO NS, and n=3 for ATF6-KO Sm; each dot represents one measurement. Data are presented as mean ± standard deviation (SD). Statistical significance was determined using an unpaired two-tailed t-test and one-way ANOVA, with p < 0.05 considered statistically significant.

### Absence of ATF6 exacerbates emphysema following repeated CS exposure

We use CS-induced COPD as a model of extreme exposure to harmful environmental toxins encountered by the lung throughout life. CS, through both active and passive inhalation, is a common environmental lung toxin that accelerates lung aging accompanied by pathological changes in the lungs (59). However, data regarding UPR activation and its role during repeated CS inhalations remain inconsistent (31), in part due to difficulty in detecting activation, which can be transient and dynamic.

We therefore investigated whether a UPR transcriptional program could be detected in mouse lungs exposed either to a single CS inhalation or to repeated twice-daily CS inhalations over six months (**Fig. 1L and Sup. Fig. 1**). After a single exposure to CS, increased transcripts encoding retrotranslocation proteins, ubiquitin ligases, and some ER quality-control factors were predominantly detected (**Sup. Fig. 1,** orange bars). Following prolonged repeated exposures, sustained induction of ATF6, IRE1, XBP1, and eIF2α mRNAs, along with the ATF6-processing proteases Mbtps1 and Mbtps2 and UPR target genes, were detected (**Sup. Fig. 1,** purple bars; (60)). These findings suggest that following a single challenge with CS, the upregulation of retrotranslocation factors and degradation enzymes is prioritized to remove irreversibly damaged proteins. During repetitive CS exposures, UPR sensors, as well as ER chaperones that restore or increase ER folding capacity and allow faster replacement of irreversibly damaged proteins, are additionally upregulated. Moreover, repetitive exposures result in increased expression of proinflammatory (TNF), proapoptotic (MAPK10, Fas, CASP8), and autophagy-related (BNIP3 and LC3) transcripts, indicating chronic activation of the UPR program that may have both protective and pro-disease roles.

We focused on comparing COPD-related pathologies in the lungs of WT and ATF6-KO mice exposed to CS for 6 months (61, 62). In WT mice, repeated CS-exposure, as expected, increased apoptotic cells in alveoli from 1.9 +/-1 cells/mm of alveolar wall to 4.7 +/-1.8 cells/mm of alveolar wall. ATF6-KO mice exhibited elevated baseline apoptosis, which remained high without further increase following CS exposure (**Fig. 1M and N**). The anti-inflammatory effect of ATF6 deficiency was partially preserved under CS exposure. The number of macrophages increased to 6.5 +/-0.25 cells/mm of alveolar wall in WT mice, but only to 4.83 +/-0.14 cells/mm of alveolar wall in ATF6-KO mice (**Fig. 1O and P**). ACL increased to 38.8 +/-1.6 μm in WT mice and to 40.0 +/-2.7 μm in ATF6-KO mice (**Fig. 1Q and R**). Moreover, ATF6 deficiency further exacerbated alveolar simplification in 20-month-old smoke-exposed mice, with ACL reaching 42.5 +/-4.6 μm and 46.7 +/-2.9 μm in WT and ATF6-KO mice, respectively (**Sup. Fig. 2A-B**). Together, these findings indicate that ATF6 deficiency accelerates alveolar simplification under both basal and smoke-exposed conditions.

### Absence of ATF6 prevents collagen deposition between small airways and blood vessels that supply them in CS-exposed mice at all ages

SAVFD represents one of the earliest pathological changes in COPD progression (23, 24, 63–65). In our model, following CS exposure, WT mice developed significant extravascular fibrosis of blood vessels that supply terminal bronchioles (mouse equivalent of small airways in humans) (**Fig. 1S** and **T**). In both 8-and 20-month-old ATF6-KO mice, SAVFD did not develop (**Fig. 1S** and **T**, right panel, and **Sup. Fig. 2C,** right panel). The divergence, where ATF6 deficiency heightens emphysema while preventing fibrotic remodeling, suggests cell-type-specific roles of ATF6 that lead to distinct pathophysiological outcomes.

### Cells that express high levels of ATF6 are present in COPD and CS-exposed lungs

To identify which cell types upregulated ATF6 expression in human and mouse lungs following exposure, we re-analyzed **s**ingle-**c**ell **RNA seq**uencing (**scRNAseq**) datasets, GSE168299 and GSE136831 (66). The human COPD atlas algorithm, built from 165,755 high-quality cells obtained from 15 control lungs not used for transplant (n = 96,303 cells analyzed) and 17 explanted lungs procured from donors with end-stage COPD undergoing transplant (with n = 69,452 cells analyzed), was used. Division of cells into major known lineages was based on the annotations and marker expression in the original Maor et al. manuscript (67) (**Sup. Fig. 3A**). The mouse scRNA dataset (66) comprised 41,104 cells obtained using an exposure protocol similar to ours, including 4 non-smoker mice (two males and two females, n = 20,246 cells analyzed) and 4 smoker mice (two males and two females, n = 20,858 cells analyzed). Cells were divided into the same major lineages as the human datasets using the same marker guidelines (**Sup Fig. 3B**).

ATF6 transcript abundance among different cell types was initially profiled using dot plot engine (68). In human COPD lungs, increased ATF6 mRNA levels were detected in **e**pithelial **P**ulmonary **N**euroendocrine **C**ells (**PNEC**), sensors cells, detecting inhaled substances, oxygen levels, and allergens; **a**lveolar **e**pithelial type **2 c**ell subclass **A** cells (**AEC2_As**), lung local progenitors of AEC1s (69–71); basal cells, function as progenitors of the ciliated and secretory cells (72, 73), and aberrant basaloid cells, cells of unclear ancestry found only in diseased lungs (**Fig. 2A**, and (74)); Elevated levels of ATF6 transcript was also detected in pericytes and **s**mooth **m**uscle **c**ells (**SMCs**), cells of stromal origins (**Fig. 2B**). Additionally, transcripts of MBTPS1 (S1P) and MBTPS2 (S2P), the proteases that process membrane-bound ATF6 into its active form, were elevated in human COPD pericytes, and MBTPS2 levels were elevated in human COPD AEC2_As (60). This provides supporting evidence for the presence of the active ATF6 processing machinery in those cell types. Within the endothelial lineage, ATF6 expression was elevated in capillary endothelial cells (**Sup. Fig. 3C**). No increase in ATF6 mRNA was detected in immune cell clusters (**Sup. Fig. 3D**).

**Figure 2.**
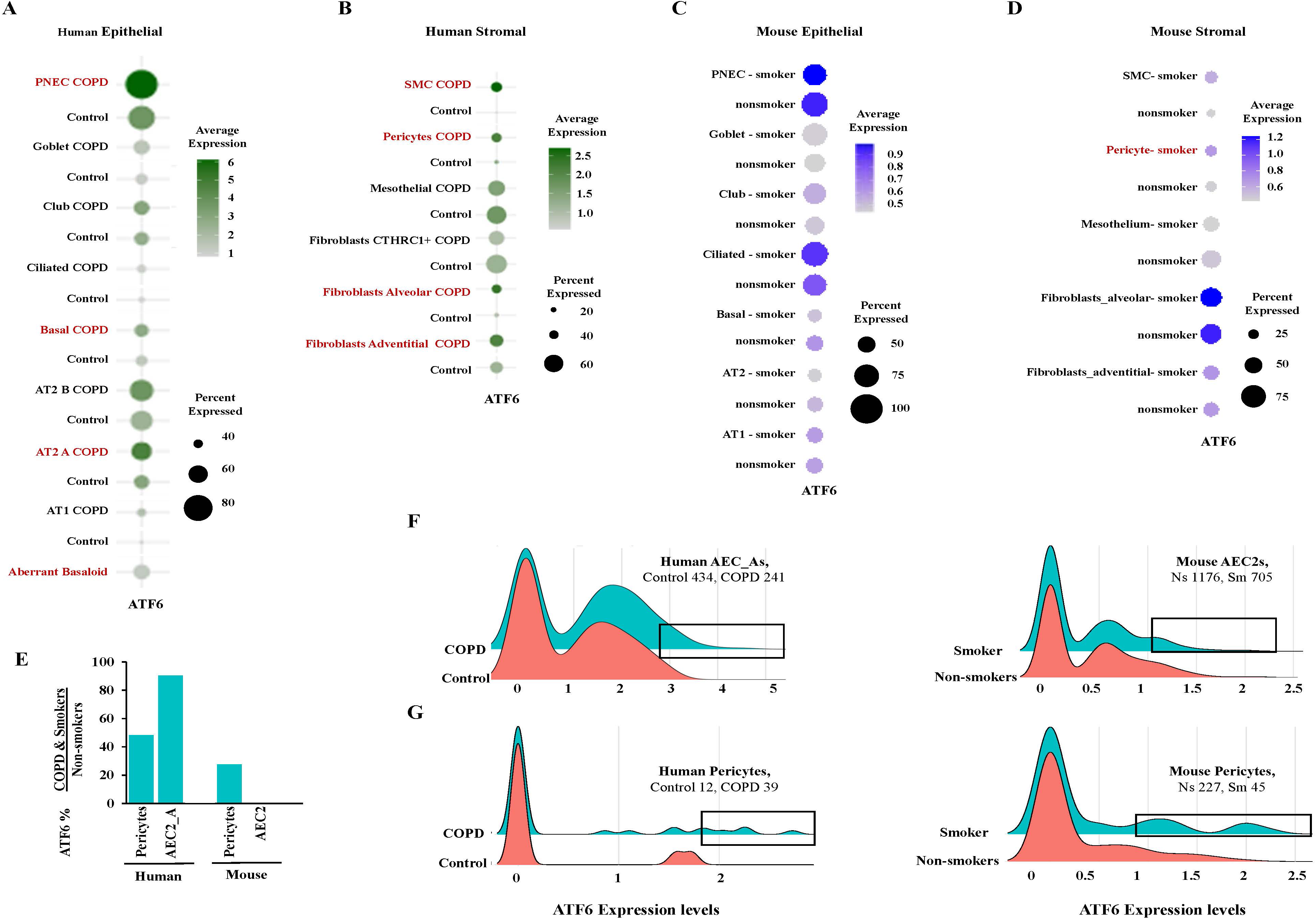
Cell types expressing elevated levels of ATF6 (ATF6^hi^) in CS-exposed mice and humans with COPD. ATF6^hi^ cells were defined as those exhibiting a ≥ 50% increase in ATF6 transcript levels relative to control counterparts. **(A - D)** ATF6 expression levels derived from scRNA-seq analysis. Dot plots demonstrate ATF6 transcript levels in human epithelial **(A)** and stromal **(B)** cells from Control and COPD lungs, and in mouse epithelial **(C)** and stromal **(D)** cells from NS and Sm lungs, respectively. Dot size reflects the percentage of ATF6-expressing cells, and color intensity represents the average normalized expression level (log-normalized counts). **(E)** Relative increase in median ATF6 expression levels across pericytes and AEC2s. **(F - G)** Ridge plots showing the distribution of ATF6 expression among AEC2s **(F)** and pericytes **(G)** in human (Control vs. COPD, left) and mouse (NS vs. Sm, right) datasets. Boxed areas contain subpopulations of cells expressing high ATF6 levels.

In mice exposed to CS, elevated ATF6 transcript was only detectable in pericytes using dot plot engine (**Fig. 2C and D**, **Sup. Fig. 3E and F**). Mouse pericytes also demonstrated 1.41-fold increase in Mbtps2 transcript. Unlike human lungs, mouse lungs did not resolve into two clear AEC2 clusters. Elevated ATF6 transcript was not detected in AEC2s.

We defined ATF6-high expressing (ATF6^hi^) cells as cell types with a ≥50% increase in ATF6 expression in human COPD lungs compared with non-COPD lungs. These ATF6^hi^ populations were pericytes and AEC2_As (**Fig. 2E)**. The relative levels of ATF6 transcript in non-exposed cells of those types were highest in AEC2_As (ATF6 relative expression level AEC2_As: 0.9261; pericytes: 0.2747), supporting previous findings on importance of ATF6 for their maintenance (75) (**Sup. Fig. 3G**). In mice when comparing bulk populations, pericytes were the only cell type classified as ATF6^hi^ in CS-exposed cells (**Fig. 2E**). This may reflect the advanced stage of the human COPD transplant specimens, whereas the mouse CS-exposure model represents earlier disease with only about 20-30% airspace enlargement detectable in smokers (76) (**Fig. 1R**). This is also suggestive that pericytes may be affected earlier in the disease, with AEC2s playing a larger role as pathology progresses.

Emphysematous areas and fibrotic airways are not distributed evenly throughout lung parenchyma, therefore, cells in affected areas may experience varying levels of UPR. To test this assumption, we analyzed those cell types using violin-ridge distribution. Heterogeneous ATF6 expression was detected within human AEC2_As, mouse AEC2s, and pericytes (**Fig. 2F-G**). We hypothesized that these ATF6^hi^ cells may represent populations localized to emphysematous areas or fibrotic niches and focused subsequent analyses on these populations.

### Transcriptomic profiling identifies mitochondrial decline in both AEC2 subpopulations and inflammatory programs in AEC2_Bs

A human AEC2_As-specific dataset of 4,089 differentially expressed genes (**DEGs**) between healthy and COPD lungs was composed and analyzed using **I**ngenuity **P**athway **A**nalysis (**IPA**). Canonical IPA revealed enrichment of differentiation-associated pathways, including MAPK, RHO, and STAT3 signaling, as well as cell cycle checkpoint and histone modification pathways involved in proliferation and differentiation (77). Downregulation of these pathways suggests impaired AEC2_As plasticity in advanced COPD. We also observed mitochondrial dysfunction, marked by downregulation of genes involved in mitochondrial energy metabolism (**Fig. 3A**). Given that mitochondrial integrity is essential for maintaining AEC2 stemness (78, 79), these findings suggest dysfunction of human lung parenchymal progenitors, AEC2_As, consistent with previous research (67).

**Figure 3.**
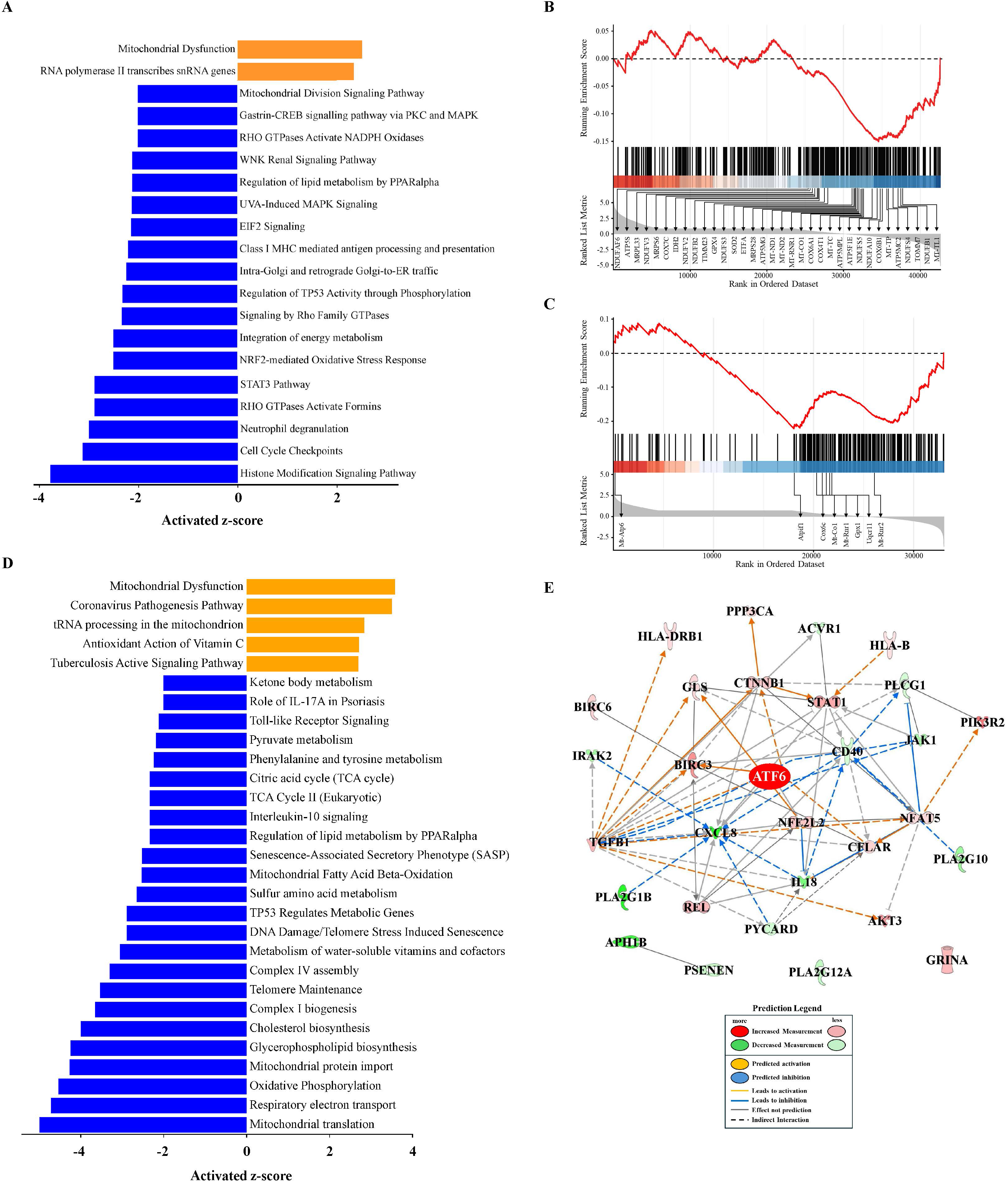
OXPHOS decline is a prominent feature of AEC2s in COPD patients, with only subtype AEC2_Bs being pro-inflammatory. (**A - C**) IPA of DEGS between healthy and COPD human AEC2_As. (**A**) The top significantly enriched canonical pathways identified by IPA in AEC2_As. Orange indicates significantly upregulated genes, blue indicates significantly downregulated genes among DEGs between controls and COPD; z-scores are plotted at the x-axis. (**B - C**) GSEA plots demonstrate downregulation of the mitochondrial-encoded and dysregulation of nuclear-encoded ETC subunits in human AEC2_As (**B**) and mouse AEC2s (**C**); genes with statistically significant expression changes are annotated. (**D - E**) IPA of DEGS between healthy and COPD human AEC2_Bs. **(D)** The top significantly enriched canonical pathways identified by IPA in DEGs from AEC2_Bs between controls and COPD. Orange indicates significantly upregulated genes, and blue significantly downregulated genes among AEC2_B DEGs; z-scores are plotted at the x-axis. **(E)** IPA-predicted gene interaction network created from the overlap between AEC2_B DEGs, GO Inflammatory Response set (GO:0006954), and ATF6. Pathway activation status was inferred using the IPA activation z-score algorithm.

To further characterize mitochondrial bioenergetics, we categorized mitochondrial DEGs as nuclear-encoded or mtDNA-encoded electron transport chain (ETC) subunits. In human AEC2_As, all mtDNA-encoded genes were significantly downregulated, whereas expression of nuclear-encoded genes for ETC subunits was dysregulated (**Sup. Table sheet 7**). **G**ene **S**et **E**nrichment **A**nalysis (**GSEA**) of the global AEC2_As transcriptome confirmed negative enrichment of ETC subunit genes (**Fig. 3B**). Although the **N**ormalized **E**nrichment **S**core did not reach statistical significance (**NES** = -0.15), individual gene analysis revealed consistent and significant downregulation of all mtDNA-encoded genes (e.g., MT-CO1, Fold change: 0.88; **Sup Table 1, Sheet 7**). We interpreted pattern as evidence that mtDNA-encoded ETC subunits are affected first, with nuclear-encoded ETC subunits becoming dysregulated as a possible compensatory response to chronic mitochondrial stress (80, 81). Same trend was found in mitochondrial of total Human AEC2s (**Sup. Fig. 4A (**GO enrichment**) and B (**IPA core analysis)). Limited number of 78 DEGs were identified in mouse AEC2s (**Sup. Table 1, sheet 8**). Those were enriched for terms such as protein refolding, chaperone-mediated protein folding and regulation of proteolysis. Due to the lack of identifiable AEC2_A populations in mice (67), we performed GSEA for ETC subunit genes on total mouse AEC2s. This analysis revealed a negative enrichment of ETC subunit genes, indicating a systematic downregulation of **ox**idative **phos**phorylation (**OXPHOS**) (**Fig. 3C**). Notably, only four mtDNA-encoded genes and four nuclear genes encoding mitochondrial proteins were significantly downregulated in smokers’ mice (**Sup. Table 1, Sheet 9**). Among these was Cox6c, a known early marker of mitochondrial decline (82), supporting the interpretation that the mouse model represents an earlier disease stage than the analyzed human COPD specimens.

Analysis of the AEC2_As subpopulation did not identify inflammation-related pathways, whereas analysis of total AEC2s showed enrichment for inflammation-related pathways (**Sup. Fig. 4A (**GO enrichment**) and B (**IPA core analysis)).

To determine whether the AEC2_Bs subpopulation, rather than AEC2_As, contributes to inflammation, a human AEC2_Bs-specific dataset of 7,796 DEGs was analyzed using IPA (**Fig. 3D**). Pathways associated with inflammation, mitochondrial dysfunction, metabolic and biosynthetic processes were enriched, consistent with previously reported AEC2_Bs features (67). Together, these results indicate distinct responses of AEC2 subpopulations to CS exposure. AEC2_As showed predominantly mitochondrial dysfunction and impaired differentiation programs, whereas AEC2_Bs additionally displayed metabolic dysregulation, and inflammation.

We next used IPA regulatory network prediction to assess whether ATF6 could contribute to these pathways. AEC2_Bs DEGs were cross-referenced with the GO:0006954 “Inflammatory Response” gene set, which predicted ATF6 as a possible upstream regulator of 26 of 30 dysregulated inflammation-related mediators in AEC2_Bs. (**Fig. 3E**). No predicted regulatory network connecting ATF6 to ETC subunit genes was identified by IPA or GO analysis in either human or mouse AEC2s.

### Transcriptomic profiling links ATF6 to vascular fibrotic remodeling in human and mouse pericytes

Human and mouse pericyte-specific DEGs were identified by comparing healthy and COPD human datasets, and sham and CS-exposed mouse datasets, respectively (**Sup. Table 1, Sheets 10 and 11**). GO enrichment analysis of mouse pericyte DEGs (**Sup. Fig. 5A**) demonstrated enrichment of ECM, vascular development, and ER stress response signatures in CS-exposed mouse pericytes.

In human pericytes, DEG analysis (**Sup. Fig. 5B**) additionally revealed enrichment of pathways involved in protein regulation and turnover, including defects in protein translation, folding, and trafficking. This may indicate severely compromised proteostasis in human pericytes, consistent with the interpretation that the human COPD datasets represent a more advanced disease stage than the mouse model.

Given that pericytes showed increased ATF6 levels in both human COPD lungs and CS-exposed mouse lungs (**Fig. 2E**), we identified cross-species common DEGs shared between the human healthy/COPD and mouse sham/CS-exposed comparisons (**Sup. Table 1, Sheets 10 and 11**).

A conserved cross-species pericyte program, consisting of 251 upregulated and 202 downregulated DEGs, was generated (**Fig. 4A**) and profiled using IPA (**Fig. 4B**). Pulmonary fibrosis, ECM organization, PDGF and TGFβ-associated signaling, wound healing, mitochondrial biogenesis, and chronic pro-inflammatory IL-23 pathways were upregulated, whereas ribosomal RNA processing, maturation and quality-control pathways were among the major downregulated pathways. This suggests that pericytes engage robust ECM remodeling and profibrotic transcriptional programs. We intersected these 453 pericyte DEGs with the GO term “extracellular matrix” (GO:0031012) and identified 16 overlapping genes (**Fig. 4C**). These 16 ECM-related DEGs formed an ATF6-ECM regulatory network, which predicted that ATF6 directly or indirectly regulates 15 of the 16 ECM-associated genes, including collagens, TIMP3, and TGFβ-related mediators. Together, these scRNAseq data position ATF6 as a central upstream organizer of pericyte-driven ECM remodeling.

**Figure 4.**
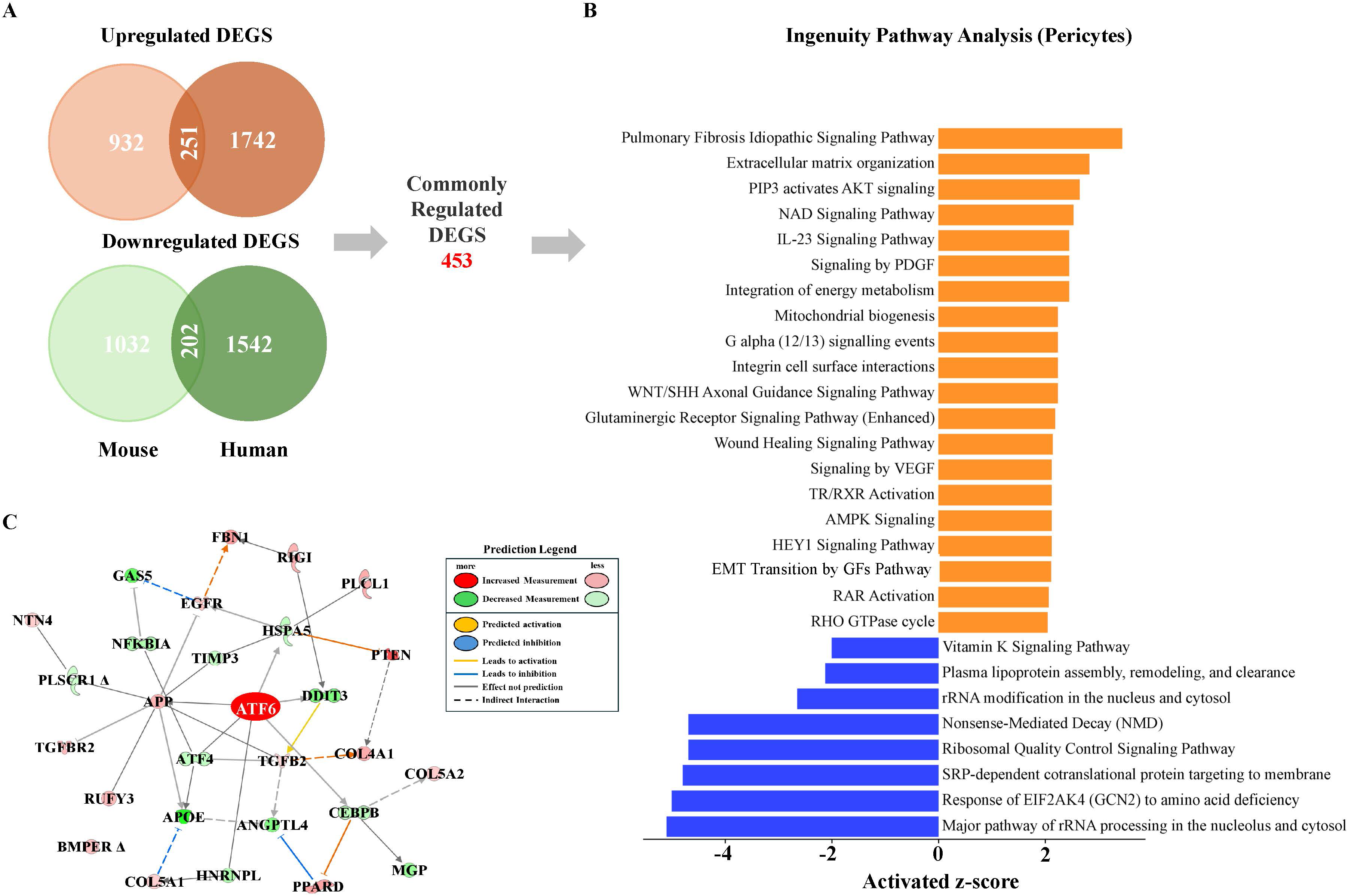
DEGs and predicted pathway analyses link ATF6 to ECM remodeling in human and mouse pericytes. IPA of pericytes transcriptomic datasets from human and mouse DEGs obtained by comparing gene expression in healthy/COPD patient and sham/smoke-exposed mice groups. (**A)** Venn diagrams demonstrate overlap of up-and downregulated human and mouse DEGs; the intersections define the similarly regulated human and mouse pericyte DEGs (n = 453). **(B)** The top significantly enriched canonical pathways identified by IPA among DEGs commonly changed between human and mouse pericytes. Orange indicates significantly upregulated pathways, and blue indicates significantly downregulated pathways; z-scores are plotted at the x-axis. **(C)** ATF6-centered ECM pathway network constructed using IPA from the overlap between the 453 shared DEGs depicted in (A) and the GO Extracellular Matrix pathway (GO:0031012). The 16 overlapping genes used to build the network are COL5A1, MGP, BMPER, COL5A2, FBN1, TIMP3, NTN4, COL4A1, ANGPTL4, RUFY3, TGFB2, APOE, PLSCR1, PLCL1, GAS5 and TGFBR2. Pathway activation status and causal relationships were inferred using the IPA activation z-score algorithm.

### Generation of multiplex panel to investigate spatial aspects of lung aging and SAVFD

Changes in mRNA levels do not necessarily translate into changes in levels of the corresponding protein (83). This discrepancy is amplified during stress, such as aging and CS-exposure, where multiple mRNAs are not efficiently translated into proteins. To validate transcriptomic findings identified in AEC2s and pericytes and predicted ATF6 regulatory roles identified via IPAs, multiplexed protein imaging panel was constructed and analyzed via the Hyperion **I**mage **M**ass **C**ytometry (**IMC**). A panel of 20 markers (**Table 1**) was designed based on reanalyzed scRNA-seq DEGs, our research (**Sup. Fig. 1,** (70)), and literature (34, 95). Targets included: cell-type specific markers; prominent profibrotic molecules; downstream ATF6 targets (ER chaperones like PDIA1, whose malfunction is linked to smoking (34, 95) and function is important for collagen processing (96), autophagy marker BNIP3, (**Sup. Fig. 1**) (98–100); senescence marker p16 (68, 101); and OXPHOS markers. Analysis of OXPHOS was further evaluated using a mitochondrial bioenergetic score calculated as the combined intensity of MT-CO1, NDUFb8, and Cox6c normalized to mitochondrial mass, estimated by VDAC1/3 (102, 103). Assuming that ATF6^hi^ cells may represent populations localized to emphysematous areas or fibrotic niches, our subsequent spatial analysis focused on profiling cells of interest localized in affected areas.

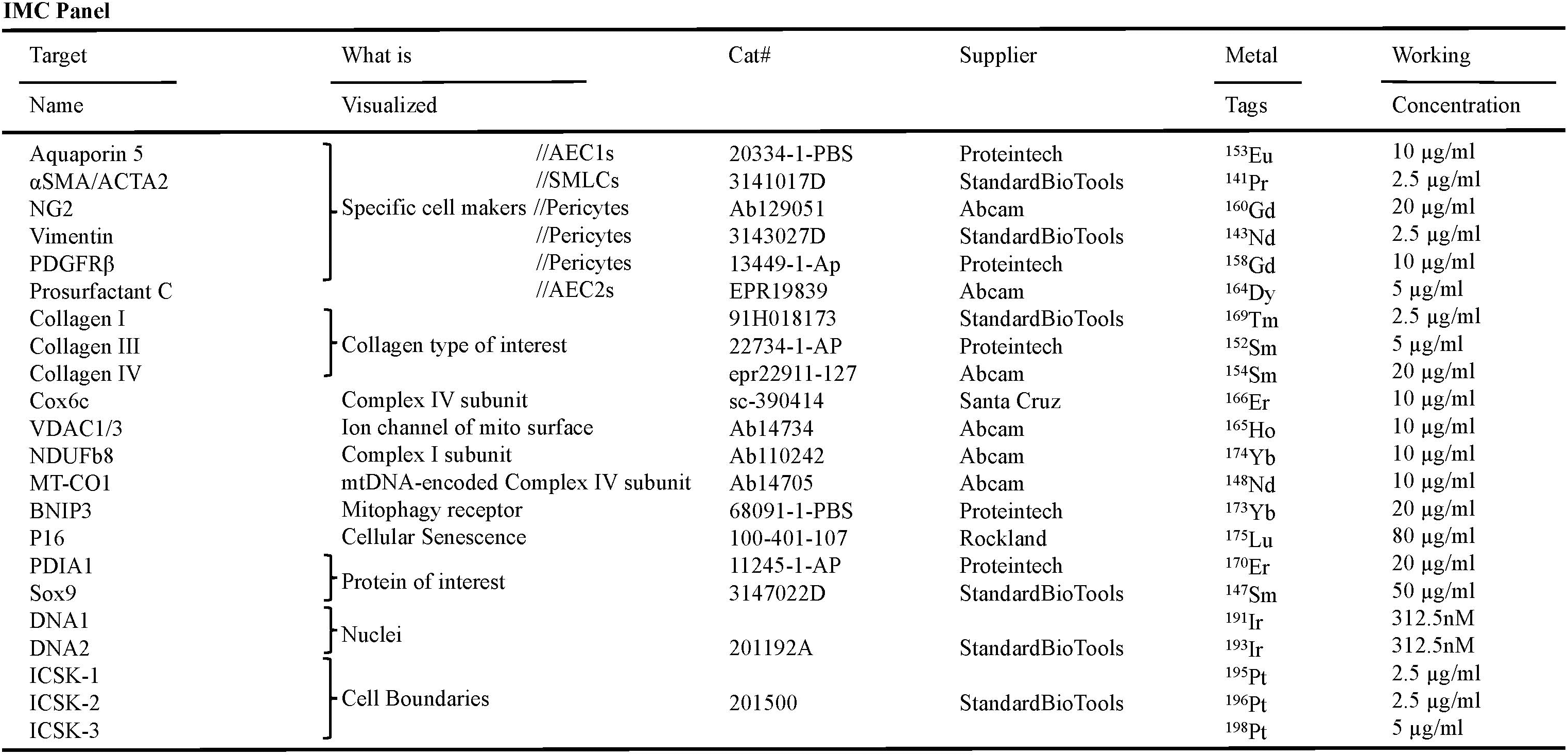

### ATF6 is required to maintain bioenergetic capacity and mitochondrial health in AEC2 progenitor subpopulations

We compared AEC2s from 8-and 20-month-old WT and ATF6-KO mice with or without CS exposure (59) (**Fig. 5A and C**). **Pr**o**s**urfactant **C**-positive (**PrSPC+)** cells, with DNA and cell plasma membrane markers defining individual cells and their boundaries, were considered AEC2s. The overall abundance of AEC2s per tissue area showed no significant differences across genotypes or exposure conditions of 8-month-old mice (**Fig. 5B**). However, smoke-exposed 20-month-old mice exhibited a significant loss of total AEC2s, and this reduction was more pronounced in ATF6-KO lungs (**Fig. 5D**).

**Figure 5.**
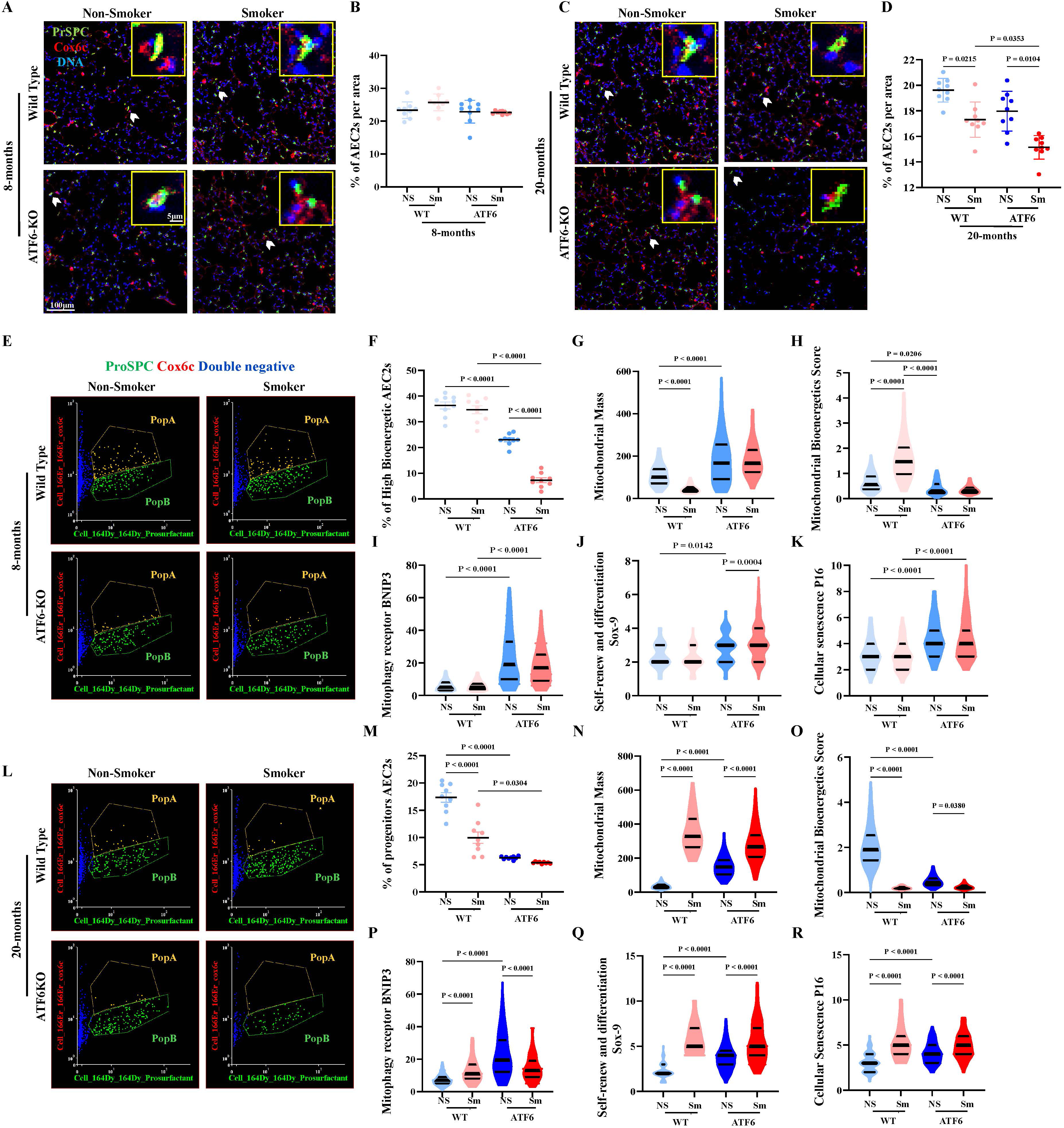
ATF6 deficiency causes age-related progressive decline in the numbers of AEC2 subpopulation with high bioenergetic capacity, which is accompanied by increased mitophagy, aberrant repair and cellular senescence. Mouse lung parenchyma from 8-and 20-month-old WT and ATF6-KO NS and Sm mice were profiled by IMC using the 20-marker panel detailed in Materials & Methods. Representative images of lung parenchyma in 8-month-old (**A**) and 20-month-old mice (**C**). Shown are PrSPC in green (^164^Dy; AEC2 marker), the mitochondrial electron transport complex IV subunit Cox6c in red (¹⁶⁶Er), and nuclear DNA in blue (^191^Ir and ^193^Ir). Inserts show enlarged arrow-marked areas where PrSPC^+^ AEC2s expressing Cox6c can be detected. Quantification of AEC2 abundance, expressed as the percentage of AEC2s per **r**egion **o**f **i**nterest (**ROI**; 500 μm × 500 μm), in 8-month-old (**B**) and 20-month-old (**D**). (**E - R**) Post-segmentation analysis of PrSPC and Cox6c expression levels in 8-month-old mice **(E - K)** and 20-month-old **(L - R)** mice. AEC2s were subdivided into population A (PopA), which expresses high levels of Cox6c and lower levels of PrSPC, and population B (PopB), which expresses lower levels of Cox6c and higher levels of PrSPC. Please note the low representation of PopA cells in ATF6-KO lung groups and in 20-month-old WT smoker lungs. PopA was further analyzed in 8-month-old **(E - K)** and 20-month-old **(L - R)** mice for the following markers. (**F, & M**) Percentage of PopA cells (high-Cox6c-bioenergetic AEC2s); **(G,** & **N)** Mitochondrial mass evaluated by VDAC levels (¹⁶⁵Ho); (**H, & O**) The mitochondrial bioenergetic score, evaluated by combined expression of two nuclear-encoded ETC subunits and one mtDNA-encoded ETC subunit per mitochondrial mass (MT-CO1 (¹⁴⁸Nd), NDUFb8 (¹⁷⁴Yb), Cox6c (^166^Er)); (**I, & P**) Mitophagy evaluated by levels of the mitophagy receptor BNIP3 (¹⁷³Yb), the transcript of which was found upregulated in CS-exposed mice (**Sup. Fig. 1**); **(J, & Q)** Aberrant self-renewal of adult AEC2s after damage evaluated by Sox9 (¹⁴⁹Sm) levels; **(K, & R)** Cellular senescence evaluated by levels of P16 (¹⁷⁵Lu). Each violin plot represents analysis of >1,000 segmented single AEC2s. Data are presented as mean ± SD. For B and D, each dot represents one region analyzed; F – K and M – R, each dot represents one individual cells. (8-month-old: WT n=3 for NS n=3 for Sm; ATF6-KO n=5 for NS and n=5 for Sm; 20-month-old: n=3 for all groups, each with technical triplicates, 3 ROIs per mouse) and were analyzed using one-way ANOVA followed by Tukey’s post hoc multiple-comparison test; p < 0.05 was considered statistically significant.

Single-cell analysis after segmentation identified two potential AEC2 subpopulations: one with low PrSPC levels (relative intensity of 6.26; PopA) and one with high PrSPC levels (intensity of 15.39; PopB) (**Fig. 5E and L**).

We further investigated ETC subunit levels and found that a subpopulation of AEC2s expressed higher levels of Cox6c (PopA relative intensity = 14.09, PopB relative intensity =9.08). Although, the AEC2_A subpopulation was not clearly identified in mice by RNAseq, we interpreted PopA from our proteomic analysis as stem-like AEC2s and PopB as more mature surfactant-producing AEC2s (AEC2_Bs). The high-Cox6c AEC2 subpopulation was reduced in ATF6-KO sham mice, with a further decline in ATF6-KO smoke-exposed lungs (**Fig. 5F**, PopA). High-Cox6c AEC2s were further depleted in CS-exposed 20-month-old WT and exposed and sham ATF6-KO mice (**Fig. 5M**), with those AEC2s being in a very low abundance in sham ATF6-KO. The adaptational increase in the ER chaperone PDI in smokers also required ATF6 **(Sup. Fig. 7)**.

The reduction in Cox6c levels was accompanied by increased mitochondrial mass in PopAATF6-KO AEC2s, and in 20-month-old WT CS-exposed mice, but not in 8-month-old WT smoker mice (**Fig. 5G and N**). The combined mitochondrial bioenergetic score, derived from Cox6c, MT-CO1, and NDUFb8 normalized to VDAC, did not change in CS-exposed 8-month-old WT mice, but was markedly reduced in 8-month-old PopA of ATF6-KO AEC2s (**Fig. 5H**), and 20-month-old WT smoker-mice (**Fig. 5O**). This decline in Cox6c is followed by a decline in cumulative mitochondrial bioenergetic score, which appeared earlier in faster-aging lungs of ATF6-KO mice, and 20-month-old CS-exposed WT mice. Increase in mitochondrial mass seems to be adaptational to the decline in bioenergetic score (Compare **Fig. 5G** and **H** of WT-Sm to **Fig. 5N** and **O**, WT-Sm). In conclusion, these data indicate that PopA ATF6-KO AEC2s exhibit substantially lower mitochondrial bioenergetic capacity, with similar phenotypes emerging in aged smoke-exposed WT mice.

Mitochondrial turnover, an indicator of dysfunctional mitochondria, was evaluated by levels of the mitophagy receptor BNIP3 (**Sup. Fig. 1**), which was highly elevated in PopA of ATF6-KO AEC2s and in 20-month-old, but not 8-month-old smoker-mice (**Fig. 5I and P**). SOX9, high levels of which has been associated with aberrant self-renewal and differentiation attempt in adult AEC2s, showed a similar trend (85) (**Fig. 5J and Q)**, as well as the senescence marker P16 (**Fig. 5K and R**).

The high-PrSPC, low-cox6c subpopulation (PopB) demonstrated similar trends in mitochondrial bioenergetic score, with surfactant levels not affected (**Sup. Fig. 8).** In conclusion, our data support the idea that ATF6 is required to maintain mitochondrial function and bioenergetic capacity in AEC2s and prevent their premature exhaustion during aging and repetitive CS exposure.

### ATF6 deficiency exacerbates AEC1 loss and mitochondrial dysfunction during aging and CS exposure

AEC1s are essential for maintaining alveolar surface architecture and gas-exchange capacity. Their renewal is AEC2-dependent, and their depletion is a hallmark of alveolar simplification (86). Consistent with alveolar simplification (**Sup. Fig. 9A**, left), IMC analysis revealed a substantial reduction in AEC1 abundance in aged smoke-exposed lungs, with the most pronounced loss occurring in 20-month-old ATF6-KO mice (**Sup. Fig. 9B**). Mitochondrial mass and bioenergetic scores in AEC1s showed an adaptive pattern similar to that observed in AEC2s: the composite bioenergetic score increased in 8-month-old WT smokers, and 20-month-old WT sham mice, consistent with age-and exposure-dependent compensation, but decreased in CS-exposed 20-month-old WT mice and in all ATF6-KO groups (**Sup. Fig. 9C and D**). BNIP3 was elevated in all ATF6-KO groups (**Sup. Fig. 9E**). P16 was not statistically increased in any test groups (**Sup. Fig. 9F**), suggesting that AEC1 renewal, rather than AEC1 senescence, is affected in ATF6-KO and in aged lungs. Together, these observations suggest that age-and ATF6-dependent dysfunction in progenitor compartments secondarily compromises AEC1 maintenance, contributing to alveolar simplification during aging and repetitive CS exposure.

### ATF6 is required for pericyte extravasation, and collagen 1 and 3 depositions

ATF6-KO CS-exposed mice did not develop SAVFD (**Fig. 1S and T**) and showed no apparent fibrotic remodeling in **n**on-**s**moker (**NS**) mice of all ages. scRNAseq data and IPA analysis demonstrated that pericytes in human COPD lungs and CS-exposed mouse lungs express high levels of ATF6, suggesting a role in regulating profibrotic remodeling (**Fig. 2**). Therefore, we tested the hypothesis that pericytes are the cells detectable within fibrotic lesions in smokers, and that their pro-fibrotic properties are ATF6-dependent.

Pericytes were identified as PDGFRβ^+^NG2^+^ or NG2^+^Vimentin^+^ cells (87–89). These double-positive cells were detected, as expected, in the walls of small blood vessels across all groups tested (**Fig. 6A, yellow arrows, and Sup. Fig. 10C-D**). Some cells were detectable within fibrotic lesions in CS-exposed WT mice and were positive for either PDGFRβ^+^NG2^+^ or NG2^+^Vimentin^+^, suggesting the presence of pericytes or pericyte-like cells within fibrotic niches in smokers. Analysis of scRNAseq data suggested that pericytes in human COPD lungs and CS-exposed mice express high levels of Col4a1 transcript (**Sup**. **Fig. 10A and B**). We did not detect Col4a1 deposition within fibrotic niches in WT smoker mice, and Col4a1 protein levels were very low within those cells (**Sup. Fig. 10C**). Col3a1 encodes a collagen implicated as an early collagen deposited during fibrotic lung remodeling and was among the DEGs upregulated in CS-exposed pericytes (fold change: 0.64; p-value 0.0079), followed by Col1a1, transcript of which was not detectable by scRNAseq (**Sup. Table 1, sheet 10**) (90). Col1 and Col3 accumulated in fibrotic areas of 8-month-old WT CS-exposed mice (**Fig. 6A**, enlarged areas). Extravascular PDGFRβ^+^NG2^+^ cells in WT smokers were strongly positive for both Col1 and Col3 (**Fig. 6A**, arrows in enlarged areas).

**Figure 6.**
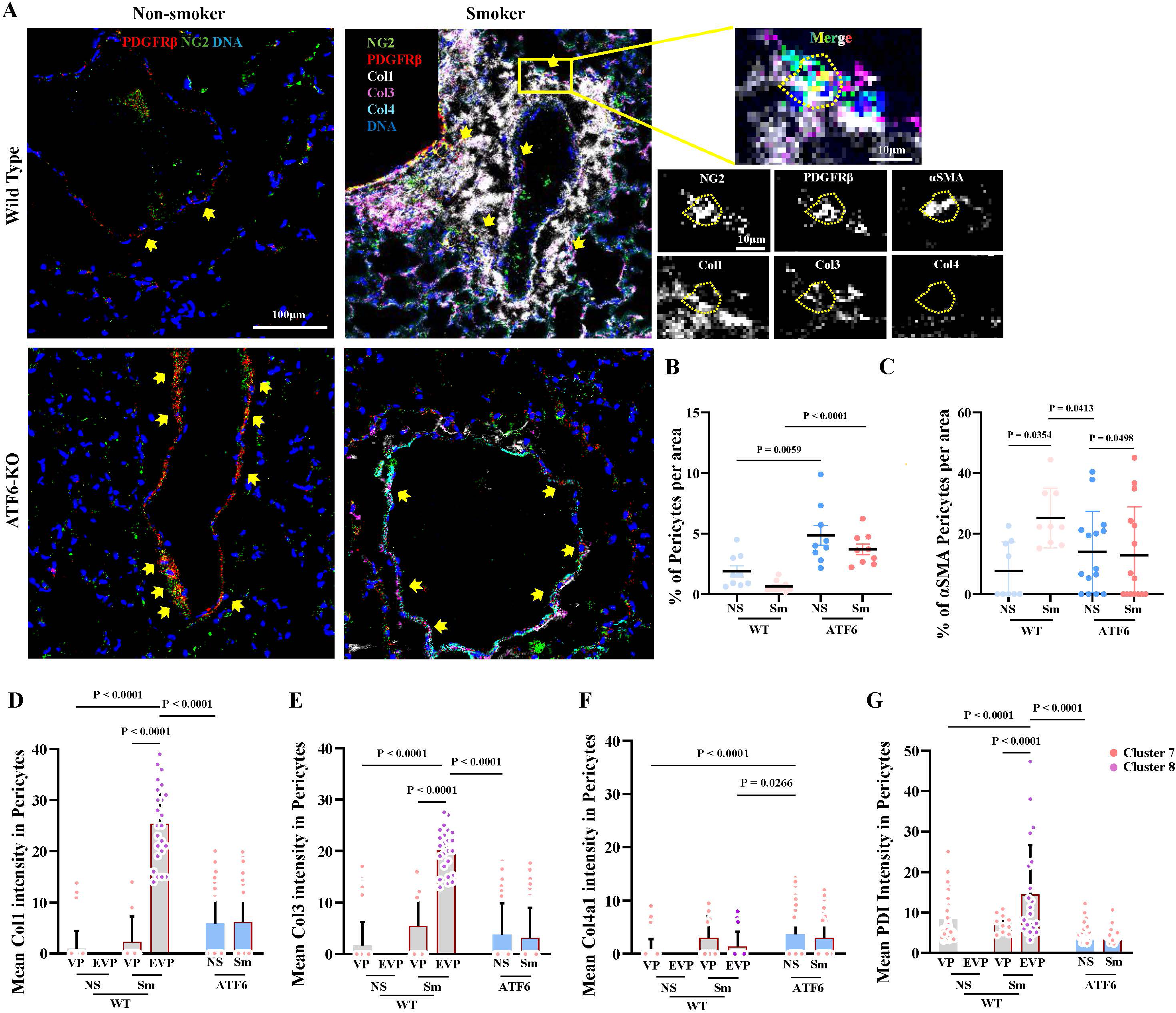
ATF6 deficiency prevents pericytes from exiting into the extravascular spaces and depositing collagen following repeated exposures to CS. Terminal bronchioles of 8-month-old WT and ATF6-KO Sm or NS mice were profiled by IMC using 20-marker panel detailed in Materials & Methods. Individual cells were identified by visualization of nuclear DNA (¹⁹¹Ir/¹⁹³Ir, blue), and cell boundaries were identified using the Cell Boundary Kit (ICSK1-3, ^195^Pt/^196^Pt/^198^Pt). **(A)** Representative images of terminal bronchioles, with pericytes identified as double NG2^+^ (¹⁶⁰Gd, green) and PDGFRβ^+^ (¹⁵⁸Gd, red) cells, lung parenchyma by Col4 (¹⁵⁴Sm), and fibrotic deposition by Col1 (¹⁶⁹Tm), and Col3 (¹⁵²Sm). Inserts display enlarged boxed areas of fibrotic airway-vascular remodeling. **(B)** Quantification of pericyte abundance, expressed as the percentage of NG2⁺PDGFRβ⁺ cells per ROI (500 × 500 μm) across experimental groups. **(C)** Quantification of pericytes re-differentiating into myofibroblasts, expressed as the percentage of NG2⁺PDGFRβ⁺αSMA⁺ cells per ROI. **(D - G)** Pericytes were subdivided into two subtypes based on their spatial localization, vascular pericytes (VP, corresponding to cluster 7 in Fig. 7) and extravascular pericytes (EVP, corresponding to cluster 8 in Fig. 7). Mean intensity of Col1 **(D)**, Col3 **(E)**, Col 4 **(F)**, and PDIA1 **(G**; ¹⁷⁰Er), an ATF6 target gene critical for assembly of procollagen trimers in the ER. Please note elevated levels of Col1, Col3, and PDIA1 only in extravascular pericytes of CS-exposed WT mice. For B and C, each dot represents one region analyzed; D – G, each dot represents an individual cells. Data are presented as mean ± SD (n=3 for WT groups and n=5 for ATF6-KO groups, each with technical triplicates, 3 ROIs per mouse) and were analyzed by one-way ANOVA followed by Tukey’s post hoc multiple-comparison test; p < 0.05 was considered statistically significant.

To characterize pericyte heterogeneity in WT mice, we manually subdivided pericytes into two sub populations, **v**ascular **p**ericytes (**VP**) and **e**xtra**v**ascular **p**ericytes (**EVP**) (**Fig. 6D-G**). Vascular PDGFRβ^+^NG2^+^ pericytes remained restricted to vessel walls and were detectable in all animal groups (**Fig. 6A**). Vascular pericytes demonstrated heterogeneous but significantly lower levels of Col1 and Col3 than extravascular pericytes (**Fig. 6D-E**). Extravascular pericytes also had higher levels of PDIA1, an ER chaperone critical for procollagen trimerization in the ER, and an ATF6 target gene (**Fig. 6G**). This is consistent with increased folding demand during pro-fibrotic remodeling. In ATF6-KO mice, extravascular pericytes were undetectable (**Fig. 6A and Sup. Fig. 10D**), and detectable pericytes did not contain high levels of Col1 and Col3 proteins nor elevated PDIA1 levels (**Fig. 6D, E** and **G**). Together, these data suggest that pericyte extravasation and subsequent collagen deposition are ATF6-dependent processes.

To determine whether pericytes exhibit energetic vulnerabilities, we quantified mitochondrial and senescence markers in the vascular and extravascular pericytes (**Sup. Fig. 10E, F-G**). No significant differences in mitochondrial bioenergetic scores or the mitophagy markers (BNIP3) were detected between WT and ATF6-KO pericytes. Nevertheless, extravascular pericytes in smoker mice were highly energetic, consistent with IPA analysis demonstrating increased energy metabolism and mitochondrial biogenesis (**Fig. 4B**).

### Transitional subpopulations of PrSPC^+^AQP5^+^ and αSMA^+^PDGFRβ^+^Col1^+^ are undetectable in ATF6-KO mouse lungs

We demonstrated (i) mitochondrial exhaustion, and low levels of AEC1s in ATF6-KO mice, with predicted dysregulation of differentiation pathways in AEC2s, (**Fig. 5 and 6 and Sup. Fig. 9**), (ii) and the presence of extravascular pericyte-like cells in WT but not in ATF6-KO CS-exposed mice (**Fig. 6 and Sup. Fig. 10**). Therefore, lineage marker expression was examined across groups using cluster analysis (**Fig. 7A-B**).

**Figure 7.**
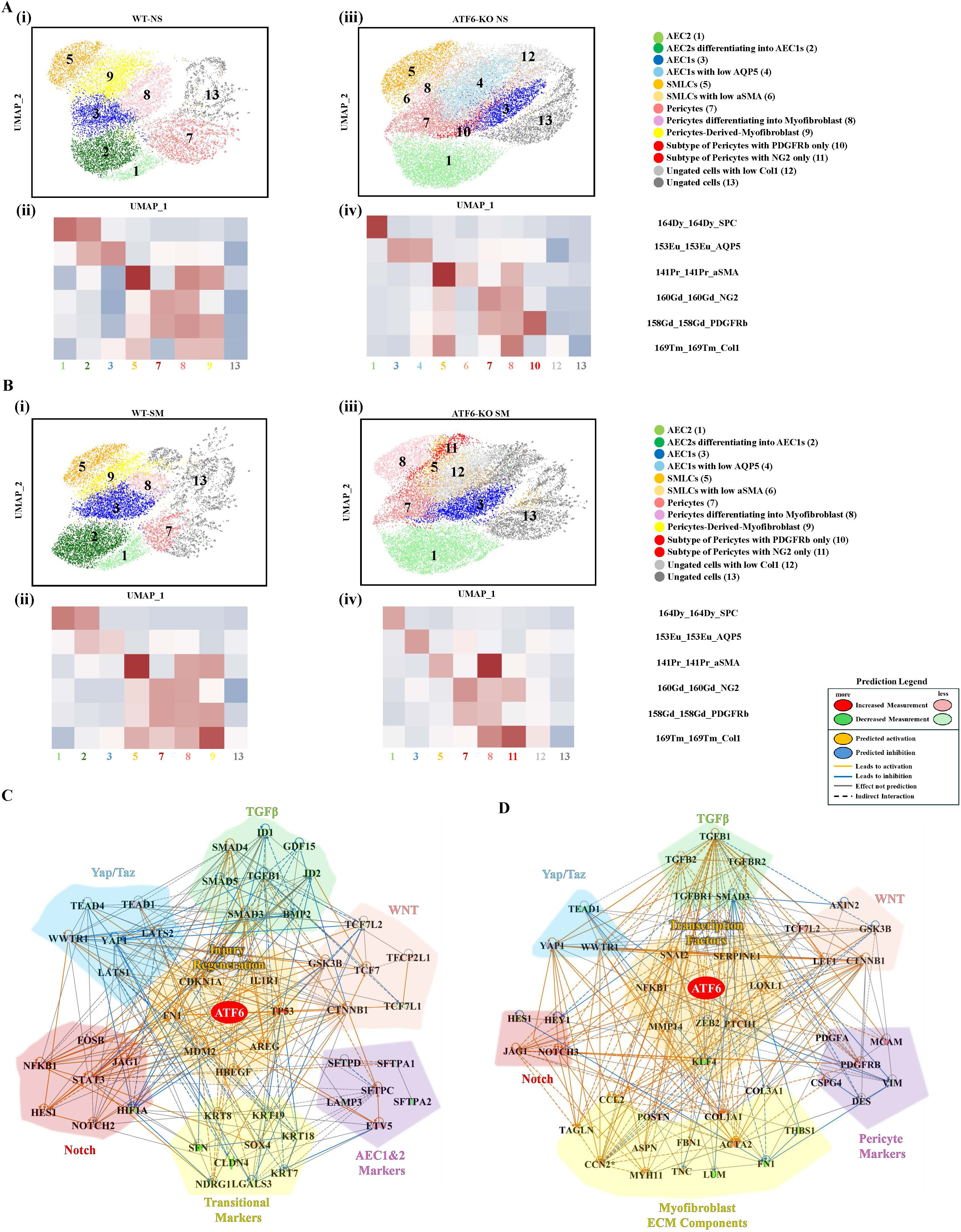
ATF6 deficiency lowers the numbers of AEC2s re-differentiating into AEC1s and abolishes the population of pericytes re-differentiating into myofibroblasts. Single-cell clustering was performed using quantitative IMC data for protein levels of cell identity markers in 8-month-old WT and ATF6-KO NS or Sm mouse lungs. Clusters are annotated based on combinatorial expression of lineage-specific markers as described in Materials & Methods. **(A, & B)** UMAPs display distinct cell-type clusters identified in WT (**i**) and ATF6-KO (**iii**) NS lungs (**A**), and Sm lungs (**B**). The corresponding heatmaps (**ii & iv**) represent the correlation of standardized marker levels across identified cell clusters. Please note that clusters 2 and 9 are not detectable in ATF6-KO lungs. Each UMAP represents analysis of >14,000 segmented single cells. (C, & D) ATF6-centered predicted differentiation pathway networks constructed from overlapping gene sets: genes required for AEC2-to-AEC1re-differentiating that overlap with DEGs between control and COPD AEC2_As (C), and genes required for the pericyte-to-myofibroblast transition that overlap with shared DEGs from human COPD and mouse CS-exposed pericytes (D). ATF6 was included in both networks. Genes associated with YAP/TAZ signaling are shown in light blue; TGFβ pathways in light green; WNT signaling in light pink; Notch signaling in light red; cell type specific markers, including AEC2_As and pericytes, are in light purple; transitional/intermediate state markers for (**C**), and myofibroblast markers and ECM components for (**D**) are in light yellow; and ATF6, injury and regeneration response genes for (**C**), and other transcription factors for (**D**) (e.g. ZEB2 for PMT driver (1) and KLF4 as a phenotypic switching regulator (2)) in the center in light orange.

In WT lungs, 8 clusters were identified: (1) AEC2s as PrSPC^+^cells; (2) AEC2s differentiating into AEC1s as double positive PrSPC^+^AQP5^+^ cells; (3) AEC1s as AQP5^+^cells; (5) SMLCs as αSMA^+^cells; (7) pericytes as double positive PDGFRβ^+^NG2^+^cells; (8) pericytes re-differentiating into myofibroblasts as triple positive PDGFRβ^+^NG2^+^αSMA^+^ cells; (9) pericyte-derived myofibroblasts as triple positive PDGFRβ^+^αSMA^+^Col1^+^cells; and (13) ungated cells. Two transitional clusters, cluster 2 (dark green), corresponding to AEC2s differentiating into AEC1s (PrSPC^+^AQP5**^+)^**, and cluster 9 (yellow), corresponding to pericyte-like cells, possibly pericyte-derived myofibroblasts αSMA^+^PDGFRβ^+^Col1^+^, were increased in CS-exposed WT groups (**Fig. 7Ai and Bi**). The increase in PrSPC^+^AQP5**^+^** cells is suggestive of accelerated AEC1 loss and the need to replace AEC1s to maintain alveolar integrity. Increase in αSMA^+^PDGFRβ^+^Col1^+^ population is suggestive of pericyte activation and exit into extravascular space, a process usually accompanied by their re-differentiate into collagen-producing myofibroblasts (**Fig. 7Ai and Bi**) (91, 92).

In ATF6-KO lungs, 11 clusters were identified: (1) AEC2s; (3) AEC1s; (4) AEC1s containing low levels of AQP5; (5) SMLCs; (6) SMLCs containing low levels of αSMA; (7) pericytes; (8) an intermediate population of pericytes re-differentiating into myofibroblasts but not mature pericyte-derived-myofibroblasts; (10) subtype of cells expressing only PDGFRβ; (11) a subtype of cells expressing NG2 only; (12) ungated cells with low Col1; and (13) ungated cells.

Clusters 2 and 9 were undetectable in ATF6-KO lungs (**Fig. 7Aiii and Biii**), suggesting that ATF6 is required for re-differentiation of AEC2s and pericytes into transitional clusters 2 and 9, respectively. Loss of cluster 2 may reflect impaired AEC2-to-AEC1 differentiation, whereas absence of cluster 9 suggests failure of pericytes to initiate myofibroblast conversion, a possible early event in fibrotic remodeling of parenchymal airways and vasculature (92).

Unexpected absence of transitional clusters in ATF6-KO lungs prompted us to put forward the hypothesis that ATF6 activity is required for successful completion of differentiation programs. To gather initial support for this hypothesis, lists of genes whose expression defines the transcriptomic programs of AEC2-to-AEC1 and pericyte-to-myofibroblast transitions were defined based on established scRNAseq atlases and lineage-tracing atlases (93–95). Those specific transcriptomic programs were compared to DEGs between Control and COPD AEC2_As and pericytes, respectively, generating sets of 64 genes that are required for AEC2-to-AEC1 transition, and 57 genes that are required for lung pericyte-to-myofibroblast transition and whose expression changes in smokers (**Sup. Table 1, sheets 12 and 13** for AEC2-to-AEC1 and pericytes-to-myofibroblast, respectively). ATF6 was included in the resulting differentiation-related sets and analyzed via IPA.

Multiple pathways that support AEC2-to-AEC1 and pericyte-to-myofibroblast differentiation were predicted to be modulated by ATF6 (**Fig. 7C and D**, respectively). For AEC2-to-AEC1 differentiation, these included YAP/TAZ of the hippo pathway, which via YAP1, WWTR1, and TEAD1/4 promote the expression of AEC1-specific genes; relative activities of TGFβ/SMAD (TGFB1, SMAD3/4/5), and BMP pathways (BMP2, ID1/2), where balance between those ensures AEC2-to-AEC1 differentiation; Notch and Wnt/β pathways that operate during different stages of the AEC2-to-AEC1 re-differentiation process; as well as induction of transitional DATP/KRT8^+^ cells (**Fig. 7C)** (96–103). Overall, analysis positioned ATF6 as an upstream regulator of 62 genes associated with AEC2-to-AEC1 transition (**Fig. 7C)**.

For pericyte-to-myofibroblast transition, ATF6 was predicted to influence Notch signaling, profibrotic ligands TGFβ1&2, and their receptors TGFBR1/2, and SMAD3, Wnt/β-catenin and Hippo/YAP pathways, and transcription factors ZEB2 and KLF4, which can activate transcription of collagen genes (**Fig. 7D)** (104–106). Crosstalk and balance between those pathways determine whether the outcome of their activity is reparative or pro-fibrotic (107, 108). Overall, analysis positioned ATF6 as an upstream regulator of 57 genes associated with pericyte-to-myofibroblast transition (**Fig. 7D)**.

Therefore, scRNAseq data support the idea that ATF6 activity is required for proper execution of cellular differentiation programs. Our data puts forward the notion that UPR is activated during cell differentiation and contributes to fine-tuning of involved pathways.

## Discussion

Decline in proteostasis networks is a central hallmark of aging and is marked by reduced ER function and efficiency (25). However, the precise contribution of the UPR to physiological and pathological lung aging remains poorly defined. Current literature is limited, largely speculative, and only now beginning to emerge conceptually, by analogy with other age-related conditions, as an important variable that may influence multiple outcomes (31). Here, we provide data from mice with targeted mutations in one of the branches of the UPR, ATF6α, that clarify its involvement in physiological lung aging and in harmful exposure-induced accelerated lung decline.

We identified two ATF6-dependent phenotypes associated with physiological aging and exposure-accelerated aging: alveolar simplification and SAVFD, respectively (**Fig. 8**). Combined analysis of scRNAseq and spatial proteomics, used to examine expression patterns and levels of proteins of interest across different lung cell types during physiological aging in WT and ATF6-deficient mice, showed that these phenotypes were driven by malfunction in specific cell types in which ATF6 transcript levels were increased in response to CS-exposure, namely AEC2s and pericytes. In AEC2s, ATF6 function was important for maintenance of stemness through preservation of mitochondrial health. In pericytes, it was important for extravasation and pathological ECM deposition outside the affected vasculature that supplies the small airways.

**Figure 8.**
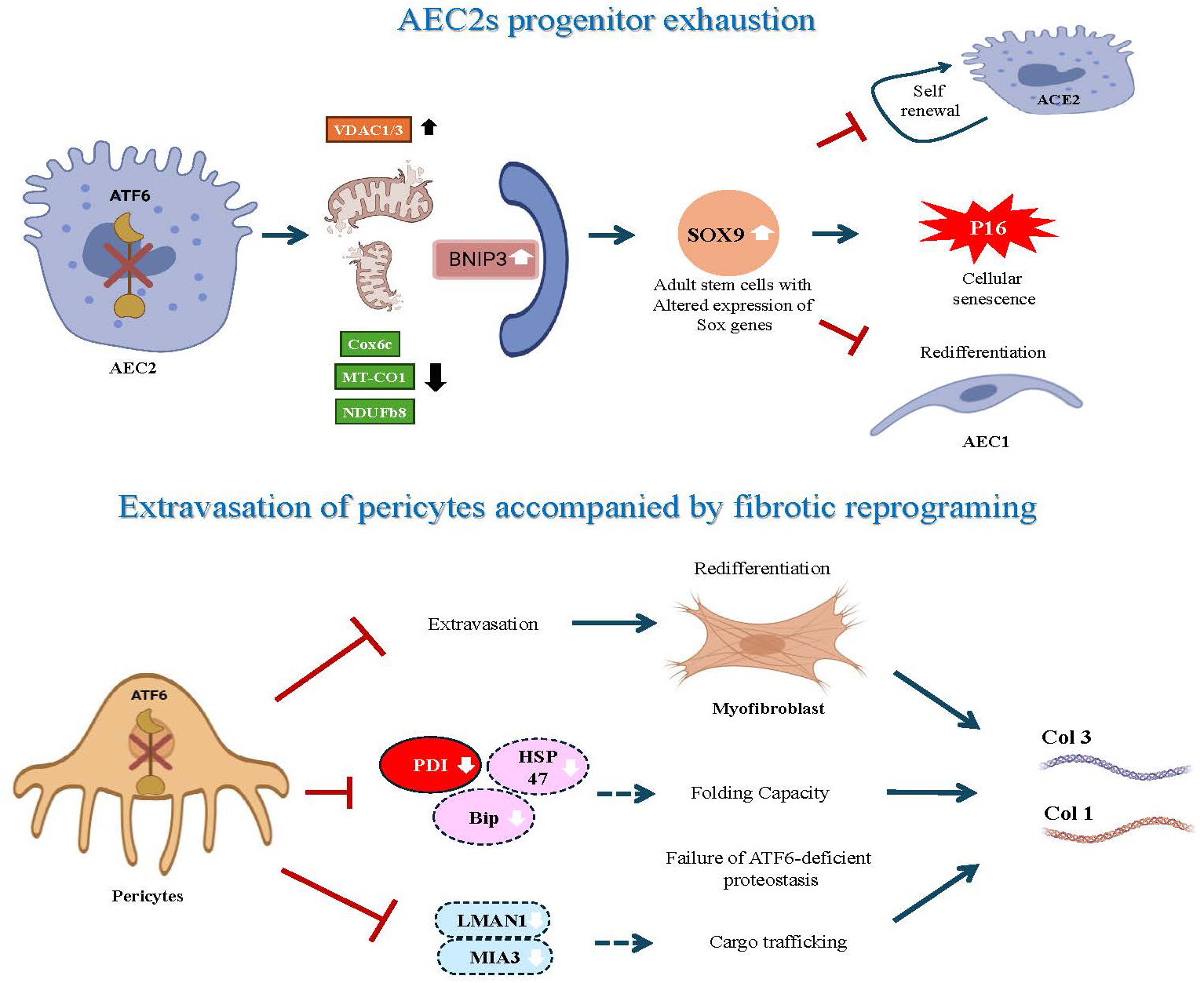
A schematic summary of the findings presented in the current report: ATF6 supports differentiation programs in AEC2s and pericytes. (**A**) Loss of ATF6 leads to age-related exhaustion of the AEC2 stem cell subpopulation. Without ATF6, high expression levels of multiple ETC subunits are not maintained. Compensatory increase in mitochondrial mass, does not improve mitochondrial bioenergetics capacity. Increased expression of BNIP3, point to the presence of dysfunctional mitochondria that is channeled for mitophagy. Elevated Sox9 level in those adult stem cells are indicative of AEC2 injury and attempted aberrant repair. Those processes culminate in impaired sell-renewal, inefficient re-differentiation into AEC1s, and cellular senescence. (**B**) Loss of ATF6 blocks vascular fibrotic remodeling around small airways driven by extravasation of pericytes and re-differentiation into myofibroblasts. ATF6-deficient pericytes showed an inability to exit into the extravascular space, re-differentiate into myofibroblasts, and deposit Col1a1 and Col3a1. Combined scRNA-seq and spatial proteomics analyses indicate that levels of ER-resident proteins required for proper collagen folding and assembly, such as PDIA1, are reduced. Supporting this observation, transcript levels of BiP and HSP47, as well as LMAN1 and MIA3, which contribute to intracellular collagen maturation and trafficking, are increased in WT mice exposed to CS. Solid lines represent pathways validated at the protein level in both WT and ATF6-deficient mice, while dashed lines indicate transcript-level observations from scRNA-seq and IPA network predictions. Created with Biorender.

Previous AEC2 analyses suggested the presence of two distinct AEC2 subpopulations with some plasticity between them: a stem cell subpopulation and a more mature surfactant-producing subpopulation (67). This division was introduced recently based on distinct gene expression signatures revealed by scRNAseq profiling. However, these two subpopulations were clearly detectable only in human, not mouse, samples (67), making their definition in mouse cells based on scRNAseq data more difficult. Our protein expression analysis detected two AEC2 subpopulations in mouse cells based on the levels of ETC subunits and prosurfactant C. Stem cell-like AEC2s (PopA) were characterized by a higher bioenergetic score and lower surfactant levels (**Fig. 5E and L**), whereas PopB of AEC2s were characterized by a lower bioenergetic score but higher surfactant C levels.

We demonstrated that mitochondrial decline in AEC2s was accelerated in the absence of ATF6. Remarkably, IPA analysis of scRNAseq expression patterns in AEC2s predicted no correlation between ATF6 and regulation of mitochondrial function (**Fig.3**), pointing to gaps in current knowledge and emphasizing the importance of our findings. In aged and smoke-exposed lungs, mitochondrial decline was characterized by decreased mRNA and protein levels of both nuclear-and mitochondrial-encoded ETC subunits per unit of mitochondrial mass, highlighting the prevalence of mitochondria with low bioenergetic scores among aged stem cell populations. A decrease in all mtDNA-encoded ETC subunits was observed in human COPD lungs from patients undergoing lung transplant, whereas a decrease in four mtDNA-encoded ETC subunits was observed in mice, which may represent an earlier stage of decline (**Sup.Table 1, Sheet 9**). Nuclear subunit transcripts were disbalanced, with decreased protein levels of complex I NDUFb8 and complex IV Cox6c subunits in both species with aging and CS-exposure. Since we can detect mitochondrial impairment in aged and exposed AEC2s, we propose use of a mitochondrial bioenergetic score described in **Fig. 5** of this manuscript and consisting of the sum of representative mtDNA-encoded and nuclear ETC components present per unit of mitochondrial mass, as determined by VDAC1/3 protein levels, as an early marker of the onset of mitochondrial decline in AEC2s.

Increased mitochondrial mass and Sox9, which may reflect an aberrant repair attempt associated with the emergence of stress-induced progenitor-like populations in the aged smoker lung, were detected in AEC2s facing energy deficit (85). BNIP3, a mitophagy receptor (111–113), was highly elevated in old and exposed mitochondria, indicating that the increase in mitochondrial mass did not result in functional improvement or healthier mitochondria. Increased P16 marker demonstrated cellular senescence in ATF6-KO AEC2s. (**Fig. 5** and **Sup**. **Fig. 8**).

All these traits were detected much earlier during the lifespan of ATF6-deficient mice (**Fig. 8**). Historically, ATF6 function has been considered protective in the majority of systems tested, although some pro-inflammatory side effects have been described (54, 55). Emerging work from M. Beers’ group describes an additional protective role for ATF6 in AEC2s. In their system, ATF6 protectively modulates UPR to mitigate pro-apoptotic IRE1α activity and downstream pro-inflammatory JNK signaling, while simultaneously promoting AEC2 proteostasis components that are essential for handling aggregated proteins (27, 75, 114).

Two recent reports suggested a link between ATF6 and ER-mitochondria Ca^2+^ homeostasis, with opposite effects on longevity depending on the experimental system and cell type examined. In human mesenchymal progenitor cells (115), ATF6 is required for maintenance of cell number and survival. The authors demonstrated an ATF6-driven transcriptomic signature specific to mesenchymal stem cells that supports their well-being through adaptive UPR. These findings support our conclusion that the ATF6 program is important in AEC2s and extend prior observations to additional stem cell types. Our data also provide supporting evidence for the idea that the ability of stem cells to mount successful differentiation programs requires ATF6. Indeed, successful redifferentiation of AEC2s into AEC1s was ATF6-dependent (**Fig. 7**). An intermediate cluster expressing markers of both AEC2s (PrSPC) and AEC1s (AQP5), at lower levels than in the single-expressing populations, was readily detectable in WT sham and CS-exposed mice but was undetectable in all ATF6-KO groups (**Fig. 5A and B**). Mitochondrial ATP production is essential for sustaining the metabolic demands of progenitor cells and differentiation programs (116–118), and disruption of ER-mitochondria cross-organelle communication could therefore have functional consequences (119).

Our previous research identified that the ER chaperone and disulfide bond isomerase PDI alters the composition of ER Mitochondria Contact Sites upon oxidation. This leads to diminished OXPHOS function due to accelerated release of cytochrome C in a threshold-dependent manner (59). Absence of ATF6 does not demonstrate adaptive increase in PDI levels **(Sup. Fig. 7B)**. Additionally, absence of ATF6 may alter levels of multiple chaperones, likely resulting in a different composition of ER-mitochondrial contact sites and additional effects on mitochondria in aging cells (120).

Another study by Burkewitz et al. (121) demonstrated the effect of ATF6 on ER-mitochondrial Ca²⁺ homeostasis and lifespan in C. elegans. In that system, the protective effect was connected to ATF6 loss and decreased levels of the ER calcium buffer calreticulin, opposite to our findings in lung parenchymal progenitor cells, further emphasizing the organismal, organ, and cell-type specificity of these outcomes. Mitochondrial bioenergetics was affected in other epithelial cell types as well, such as AEC1s. Taken together, absence of ATF6 resulted in premature alveolar simplification that was age-dependent, accelerated by harmful exposure, and accompanied by loss of AEC2s and AEC1s.

The importance of ATF6 in disease development has been overlooked in comparison to other molecules of the UPR pathway (54). Here, we provide evidence for the pathogenic importance of ATF6 signaling in lung pericytes. Our analysis revealed that one of the main pathophysiological characteristics of COPD, SAVFD, is not evident in ATF6-deficient mice in the CS-induced COPD mouse model (**Fig. 1S and T**). SAVFD develops at the intersection of small airways, the vasculature that supplies them, and alveoli, and may accelerate alveolar simplification by disturbing alveolar homeostasis and creating emphysematous areas adjacent to fibrotic lesions. SAVFD is usually not observed during physiological aging. Consistent with this, normal connective tissue around small airways and blood vessels was unaffected in ATF6-KO unexposed mice at any age tested (**Fig. 6A and Sup. Fig. 10D**).

Mining of pre-existing scRNAseq datasets identified lung pericytes as the only cell type showing a significant increase in ATF6 levels due to exposure in both mice and humans, indicating that pericyte malfunction may be an early and important factor in the disease pathology of accelerated aging (**Fig. 2**).

Pericytes are crucial mesenchymal cells surrounding microvessels and contributing to structural integrity and vascular function. Importantly, in WT mice, sham and smokers, we were able to detect pericytes located in the blood vessels that supply small airways. These pericytes did not demonstrate any pro-fibrotic signatures (**Fig. 6A**), supporting previous data (123). In contrast, profibrotic, extravascular pericyte-like cells were present only in CS-exposed WT mice within the lesions. These cells were positive for PDGFRβ^+^, NG2^+^, αSMA^+^, and produced high levels of Col1 and 3, fitting a myofibroblast profile. In agreement, the lung myofibroblast population has been shown to originate from pericytes, interstitial fibroblasts, and MSCs (123). These PDGFRβ^+^, NG2^+^, αSMA^+^ cells were undetectable in ATF6-deficient smoker-mice (**Fig. 7A and B**), suggesting that re-differentiation of pericytes into myofibroblasts requires ATF6. Thus, our data strongly suggest that ATF6 governs the adjustment and optimization of ER function required to support multiple cellular differentiation programs, which may have both protective and disease-promoting effects. It also emphasized the importance of spatial context of analyzed cells, where the information on extravascular pericytes like cells would be lost without knowledge of their localization.

A few processes required for pericyte re-differentiation into myofibroblasts may rely on ATF6-mediated ER adaptational plasticity: rapid production of motor proteins that ensure cellular movement out of the vasculature, procollagen synthesis itself, and successful re-differentiation program.

It is currently accepted that levels of major lung collagens, including types 1, 3, and 4, with Col 4 being a network-forming collagen that supports alveolar structure, while Col 1 and 3 are fibril-forming collagens (124), remain unchanged or are even increased in old versus young mouse lung. This situation changes in COPD patients due to repetitive exposure, when increased degradation of ECM proteins by **m**atrix **m**etallo**p**roteinases (**MMPs**) and neutrophil elastase occurs (124), creating a need for repair, where extravasation of lung pericytes may represent one such repair attempt.

To sustain these repair attempts, the ER of re-differentiating pericytes must possess strong ER-resident machinery for procollagen biosynthesis, post-translational modification, and triple helix formation (124). Some of ER collagen-specific chaperones BiP, Grp94, PDI, calreticulin, calnexin, are all ATF6 pathway downstream targets. Additionally, procollagen export from the ER is COPII and TANGO1/HSP47-dependent, with COPII being upregulated in smokers pericytes.

ATF6 deficiency may impair procollagen synthesis in and exit from the ER, creating heightened toxic ER overload with accumulating unfolded pro-collagen cargoes and making those cells susceptible to death. Therefore, ATF6 pathway in CS-exposed pericytes ER stress is disease-promoting, via effects on procollagen synthesis in ER.

An additional point we would like to emphasize is that scRNAseq detected increased Col4A1 and Col3A1 transcripts in CS-exposed pericytes, but not Col1A1 (**Sup**. **Fig. 10A and B**). While absence of Col1A1 transcripts can be explained by being below scRNAseq detection levels (**Fig. 2G**), the discordance between Col4A1 transcript and protein levels raises the possibility that not all transcripts are translated into proteins, especially under conditions of chronic stress. This further suggests that evaluation of both RNA and protein levels is required to fully understand diseased phenotypes, in which increased transcript levels may represent a failed adaptation/repair attempt, with mRNA unable to generate protein (83). This idea is further supported by IPA analysis of pathways demonstrating that ribosomal assembly and ribosomal subunits are downregulated in human COPD pericytes, which represent late stage of disease. Notably, mitochondrial bioenergetics and BNIP3 levels were not affected in CS-exposed lung pericytes. These cells, especially extravascular pericytes, were highly energetic (**Sup. Fig. 10E**). Absence of ATF6 had no effect on the bioenergetics of remaining/intravascular pericytes, further pointing to the cell-type specificity of ATF6 programs.

Important findings from our work lead to the conclusion that ATF6 is essential for the ability of cells to mount and maintain complex differentiation programs in epithelial AEC2s and mesenchymal pericytes. Absence of ATF6 compromised the complex and rapid cellular adaptation required during differentiation/re-differentiation reprogramming, resulting in decreased differentiation capacity. The outcomes of this are strictly cell-type specific and can compromise both beneficial processes, stemness and AEC2s → AEC1s transitions, and pathological processes, pericyte → myofibroblast transitions and adaptive responses. This cell-type specificity may reflect complex regulatory networks recently suggested by Jiménez-Beltrán and colleagues. In our case, those are formed via interaction of cell type specific transcription factors with ATF6 (**Fig. 7**) and govern physiological activities of ATF6 that extend beyond the traditional roles in UPR (52). Our work unequivocally demonstrates the importance of understanding cell-type-specific ATF6 programs in both physiological and pathological conditions. ATF6 modulators, both activators and inhibitors, have been developed. Therapeutic use of these modulators will need to be very carefully titrated and restricted to specific cell types and conditions.

## Contributions

ABP designed the study, performed multiple experiments, oversaw data analysis, and wrote the manuscript with inputs from other co-authors. XH performed IMC experiments, titrated IMC panels, designed analysis workflow, analyzed proteomic data, performed IPA and GO analyses, wrote manuscript and generated figures with inputs from other co-authors. JEB performed single cell analysis and assisted in single-cell clustering of IMH data. SJM performed initial analysis of ATF6 cell-type specific expression using COPD atlas. BIT and YB performed initial and assisted XH with IPA analyses. YB critically read the manuscript. KV, CEN, HK, CER and NAP maintained animal colony, performed smoke exposure, lung collection and analyses of the relevant lung phenotypes. NAP edited the manuscript. ASL helped with initial morphometry analysis. SDS, and YP helped with initial design, definition of lung pathological phenotypes, and critical reading of the manuscript.

## Acknowledgements

This research was supported by Aging Seed Funding Award, University at Buffalo, and American Heart Association, Transformational Award number 25TPA1481544 to ABP, National Institute of Health/ National Heart Lung and Blood Institute (NIH)/(NHLBI) grant R01HL163168 to YB. The funders had no role in study design, data collection and analysis, decision to publish or preparation of the manuscript. We would like to thank Randal J. Kaufman, the director and a professor of the Degenerative Diseases Program, Neuroscience and Aging Center at Sanford Burnham Prebys Medical Discovery Institute for a gift of ATF6α knockout mice. Jiantao Pu, Professor of Radiology and Bioengineering, Department of Radiology, University of Pittsburgh for sharing AI morphometry algorithm. Imtiaz Mohammad, Director Histology Core/Lead HistoTechnologist, and Wade Sigurdsson, Director Pathology Anatomy Sciences Research Service Center, JSMBS, UB core facility for the precise tuning and operational setup of the imaging mass cytometry system, and for the help in the use of Hyperion IMC. Special thanks to Standard BioTools team, Nina Lane, Senior Product Manager at Standard BioTools for inviting us to the service site and teaching us to use Hyperion mass cytometer, to Vinicius Motta, Senior Field Applications Scientist at Standard BioTools, for providing foundational training in data analysis methodologies, to Lindsy Rapkin, Field Applications Specialist at Standard BioTools, for critical assistance with antibody titration, channel concentration optimization, and the establishment of analytical thresholds. We also extend our gratitude to Roy Miller for technical help with immunofluorescence for antibodies validation.

## Footnotes

## Competing interests

The authors declare no competing interests.

ACL: Average Chord Length
AEC1s: Alveolar Epithelial Type 1 Cells
AEC2s: Alveolar Epithelial Type 2 Cells
AEC2_As: Human AEC2 progenitor’s subpopulation,
AEC2_Bs: Human AEC2 surfactant producing mature subpopulation
AEC2s-PopA: Mouse AEC2 subpopulation with high Cox6c expression
AEC2s-PopB: Mouse AEC2 subpopulation with lower Cox6c higher prosurfactant C expression
AQP5: Aquaporin 5
αSMA: Alpha Smooth Muscle Actin
ATF6: Active Transcription Factor 6 α
ATF6-KO: Global ATF6-Deficient
BNIP3: BCL2 Interacting Protein 3
Col1: Collagen Type I
Col1a1: Collagen Type I Alpha 1 Chain
Col3: Collagen Type III
Col3a1: Collagen Type III Alpha 1 Chain
Col4: Collagen Type IV
Col4a1: Collagen Type IV Alpha 1 Chain
COPD: Chronic Obstructive Pulmonary Disease
COPII: Coat Protein Complex II
CS: Cigarette Smoke
DEGs: Differentially Expressed Genes
ECM: Extracellular Matrix
ER: Endoplasmic Reticulum
ERSR: Endoplasmic Reticulum Stress Response
ETC: Electron Transport Chain
FDR: False Discovery Rate
FFPE: Paraffin-Embedded Formalin-Fixed
GEO: Gene Expression Omnibus
GO: Gene Ontology
GSEA: Gene Set Enrichment Analysis
HSP47: Heat Shock Protein 47
IHC: Immunohistochemistry
IMC: Imaging Mass Cytometry
IPA: Ingenuity Pathway Analysis
IRE1α: Inositol-Requiring Enzyme 1Α
KEGG: Kyoto Encyclopedia of Genes and Genomes
MT-CO1: MtDNA-Encoded Subunit of Complex IV
mtDNA: Mitochondrial DNA
NDUFb8: NADH:Ubiquinone Oxidoreductase Subunit B8
NG2: Neuron-Glial Antigen
OXPHOS: Oxidative Phosphorylation
PCA: Principal Component Analysis
PDGFRβ: Platelet-Derived Growth Factor Receptor β
PDI: Protein Disulfide Isomerase
PERK: Protein Kinase RNA-Like Endoplasmic Reticulum Kinase
PNEC: Epithelial Pulmonary Neuroendocrine Cells
PrSPC: Prosurfactant C
ROI: Regions Of Interest
S1P: Site-1 Protease
S2P: Site-2 Protease
SAVFD: Small Airways and Vasculature Fibrotic Disease
scRNAseq: Single-Cell RNA Sequencing
SMLCs: Smooth Muscle Like Cells
Sox9: SRY-box Transcription Factor 9
TANGO: Transport and Golgi Organization protein 1
TGFβ: Transforming Growth Factor β
TUNEL: Terminal Deoxynucleotidyl Transferase dUTP Nick-End Labeled
UMAP: Uniform Manifold Approximation and Projection
UPR: Unfolded Protein Response
VDAC: Voltage-Dependent Anion Channel
VE: Vascular Endothelial
VP: Vascular Pericyte
EVP: Extravascular Pericyte
WT: Wild Type

## Supplemental Figure Legends

**Supplemental Figure 1.**
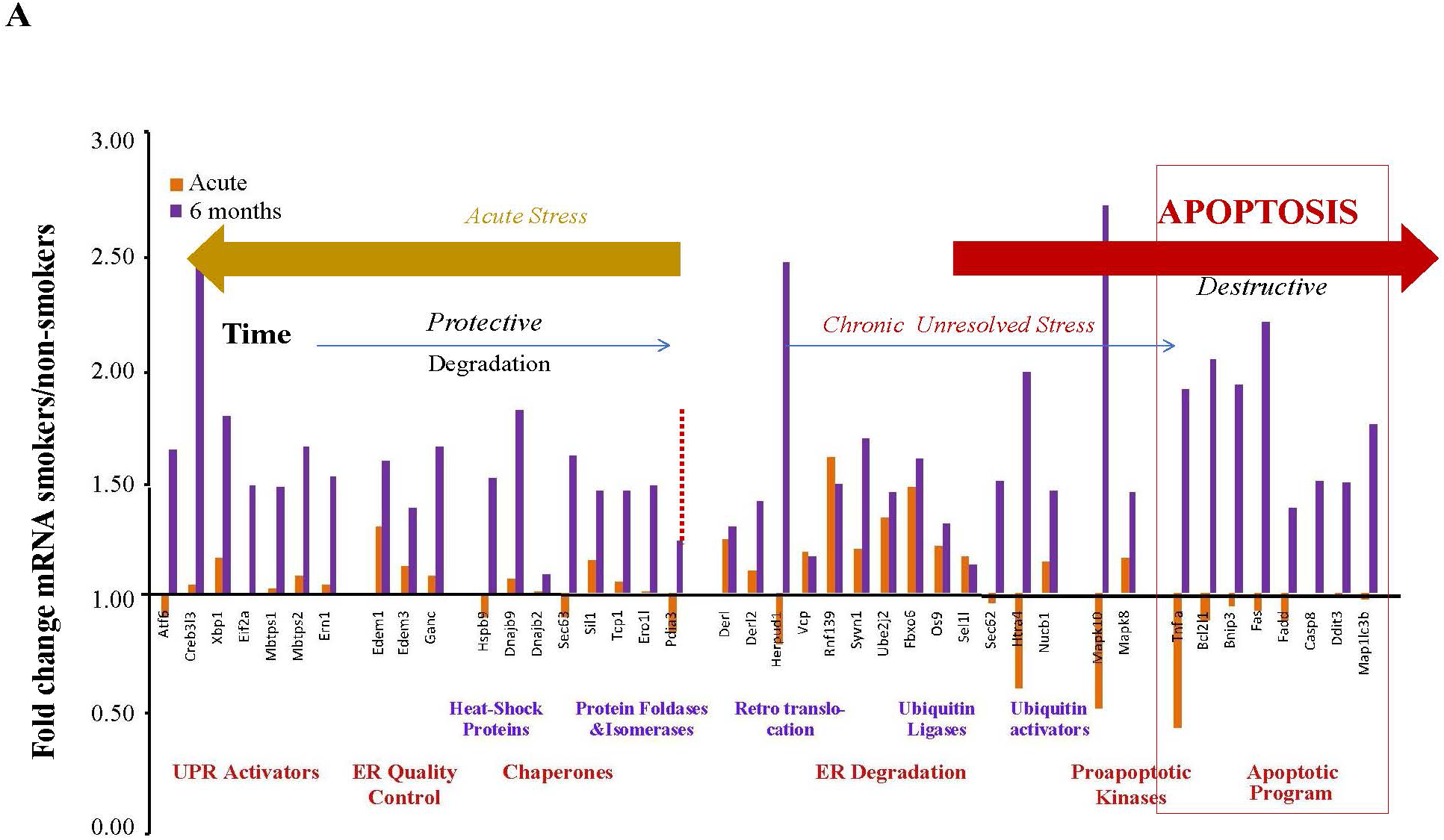
UPR program in mice subjected to single and repetitive CS exposures. 8-week-old mice were subjected to either a single CS exposure or repetitive CS inhalation twice daily for 6 months (**Fig. 1L**). RNA was extracted from total lungs 16 hours post last exposure. Three total lung lysates were pooled together for each condition and analyzed for gene expression changes using Qiagen UPR RT2 Profiler PCR Arrays. All gene expressions were normalized to GAPDH. Genes with fold changes ≥1.7-fold were grouped based on known function, where orange bars represent single CS exposure and purple bars represent repetitive CS inhalations. Gene grouping illustrates the potential shift from protective adaptive responses following single exposure to destructive components following repeated exposures.

**Supplemental Figure 2.**
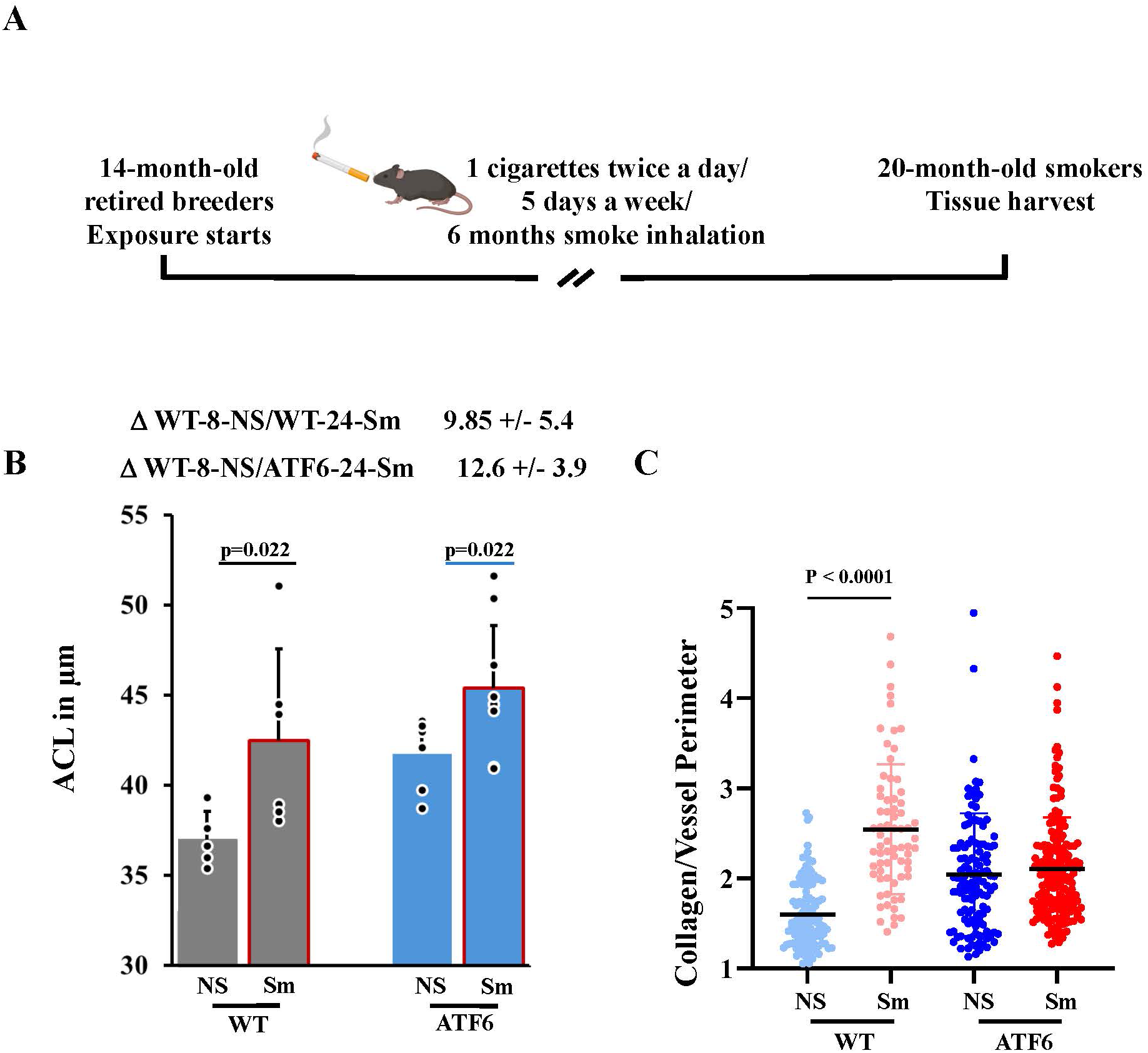
Repetitive CS exposure in aged mice accelerated alveolar simplification and vascular collagen accumulation. **(A)** Schematic representation of experimental design. **(B)** Quantification of ACL; each dot represents the average of 8-12 fields per animal (n=4 female and 4 male for WT NS, n=4 female and 2 male for WT Sm, n=3 female and 4 male for ATF6-KO NS, and n=3 female and 4 male for ATF6-KO Sm). **(C)** Collagen deposition around blood vessels that supply small airways was quantified from Masson’s Trichrome stained lung sections as the collagen/vessel perimeter ratio, with each dot representing one airway (25-30 airways per animal, n=1 female and 4 male for WT NS, n=3 female and 2 male for WT Sm, n=3 female and 4 male for ATF6-KO NS, and n=5 female and 4 male for ATF6-KO Sm). Data are presented as mean ± SD and were analyzed by one-way ANOVA with Tukey’s post hoc multiple-comparison test; p < 0.05 was considered statistically significant.

**Supplemental Figure 3.**
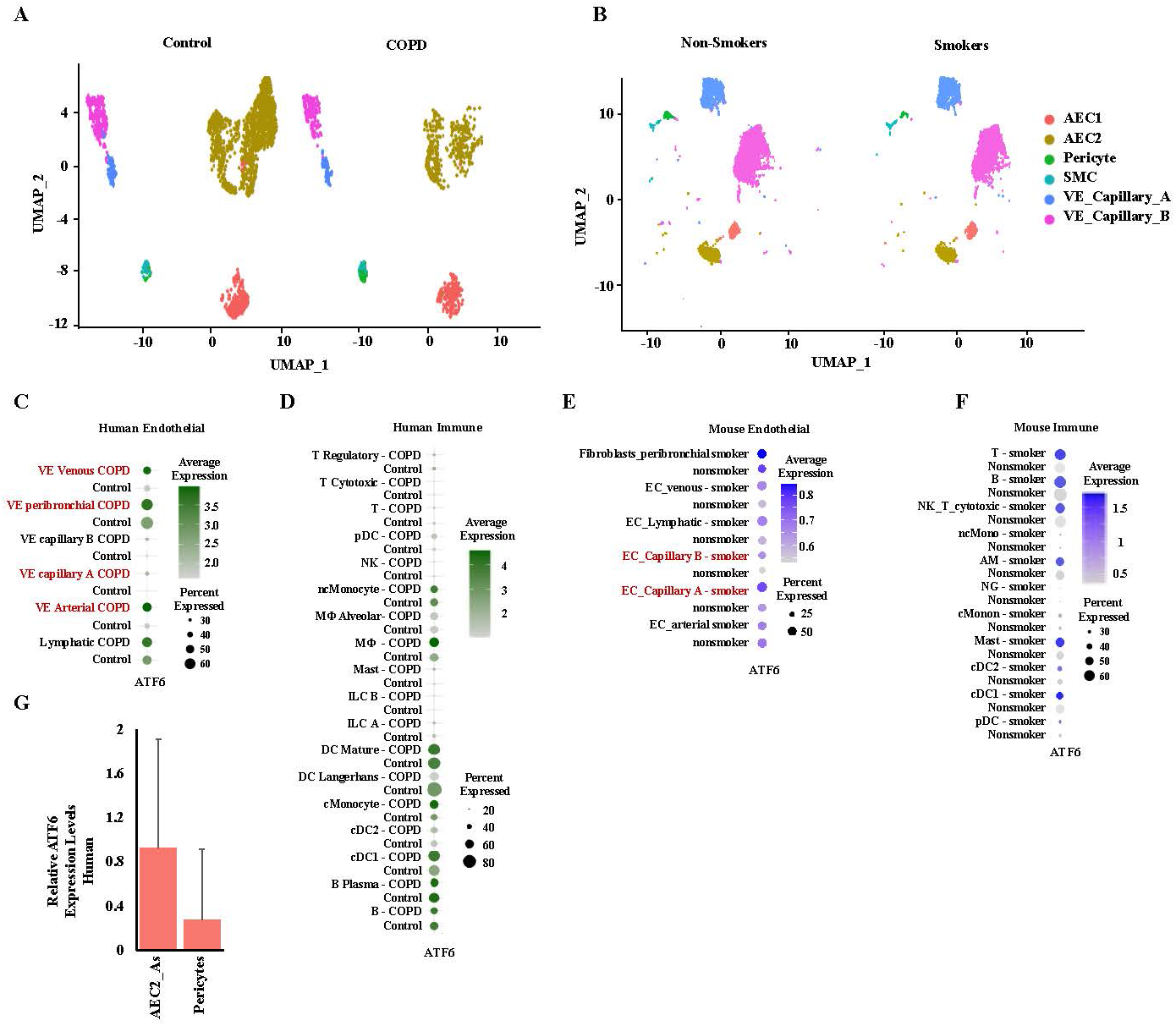
Single-cell transcriptomic visualization of ATF6 expression in human and mouse lungs. Publicly available scRNA-seq datasets from human (GSE136831) and mouse (GSE168299) lungs were analyzed using Seurat v4.1.0 to assess ATF6 transcript distribution across pulmonary cell populations. UMAP visualization of single-cell transcriptomes showing major lung cell populations, including AEC1, AEC2, pericytes, SMCs, and **v**ascular **e**ndothelial (**VE**) cells in human (**A**) and mouse (**B**) lungs. **(C - F)** Dot plots summarize normalized ATF6 expression across endothelial and immune cell subsets in human **(C - D)** and mouse **(E - F)** lungs. Dot size represents the percentage of ATF6-expressing cells, and color intensity indicates the average normalized expression level (log-normalized counts). (**G**) Bar plot showing relative ATF6 expression levels across human AEC2_As and pericytes.

**Supplemental Figure 4.**
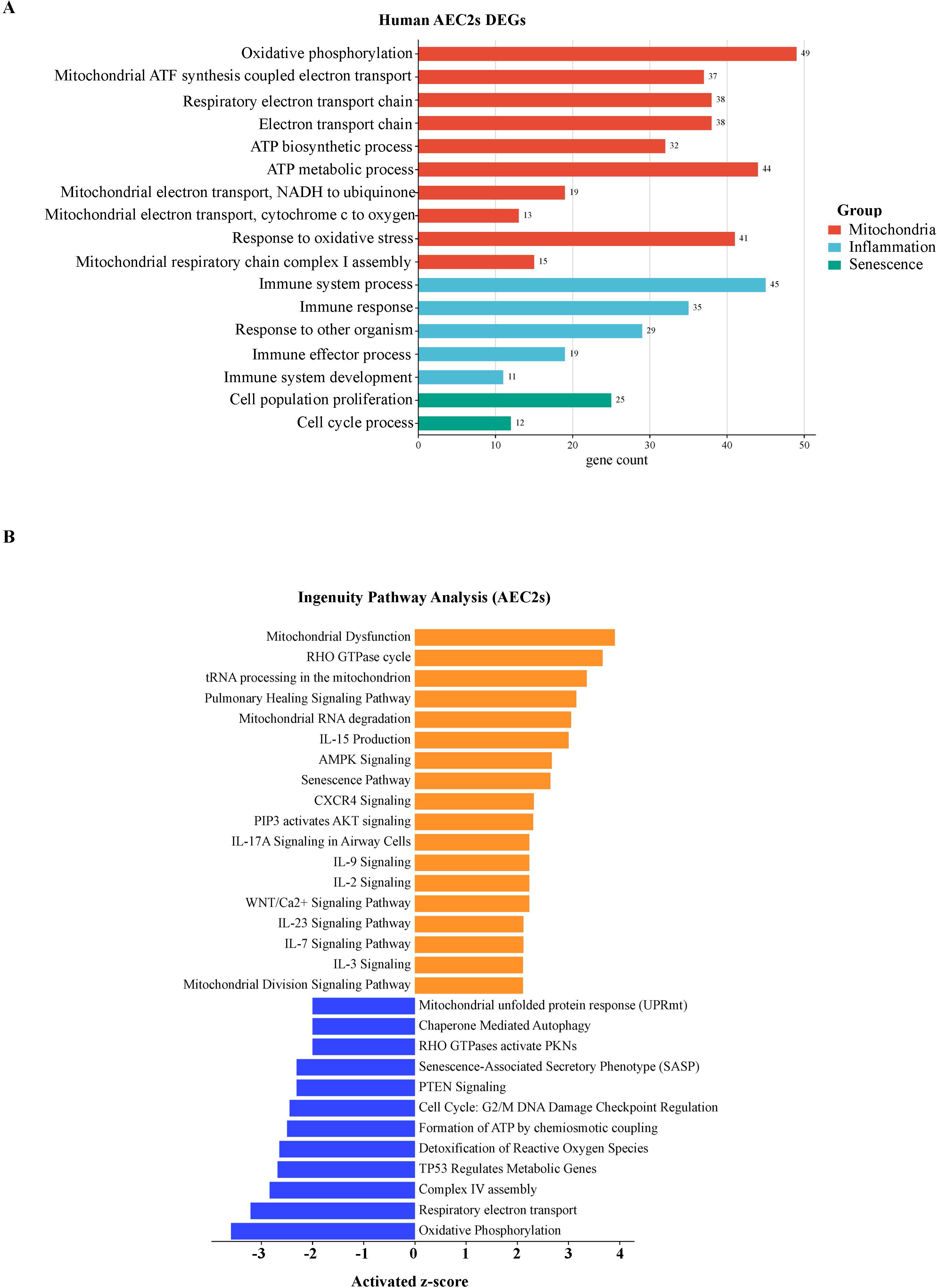
GO enrichment and IPA core analysis of human AEC2s. **(A)** DEGS between healthy and AEC2s from COPD patients were analyzed. The bar chart displays the top enriched terms categorized by functional groups: Mitochondria (orange), Inflammation (light blue), and Senescence (green). The length of each bar and the adjacent number represent the gene count associated with each specific GO term. (**B**) The top significantly enriched canonical pathways identified by IPA in AEC2. Orange indicates significantly upregulated genes, and blue significantly downregulated genes from the experimental dataset; z-scores are plotted on the x-axis.

**Supplemental Figure 5.**
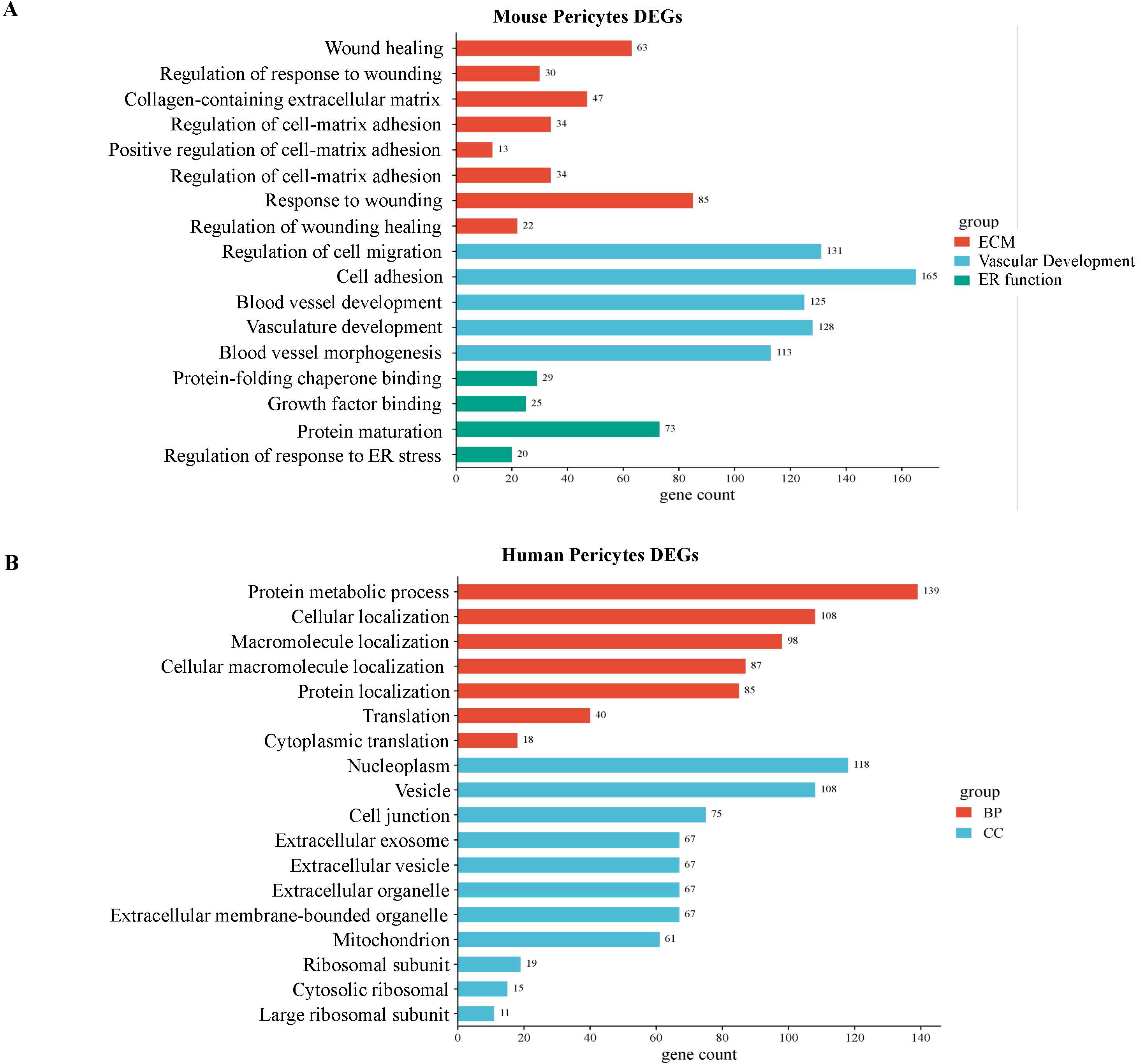
GO enrichment analysis of human and mouse lung pericytes. (**A)** Bar chart showing enriched GO terms among DEGs between sham and smoke-exposed mouse pericyte group. Terms are categorized into functional groups: Extracellular Matrix (ECM, orange), Vascular Development (light blue), and ER function (green). The x-axis represents the gene count associated with each GO term. **(B)** Bar chart showing enriched GO terms among DEGs between healthy human pericytes and once from COPD patients. Terms are categorized by GO domain: Biological Process (BP, orange) and Cellular Component (CC, light blue). The x-axis represents the gene count.

**Supplemental Figure 6.**
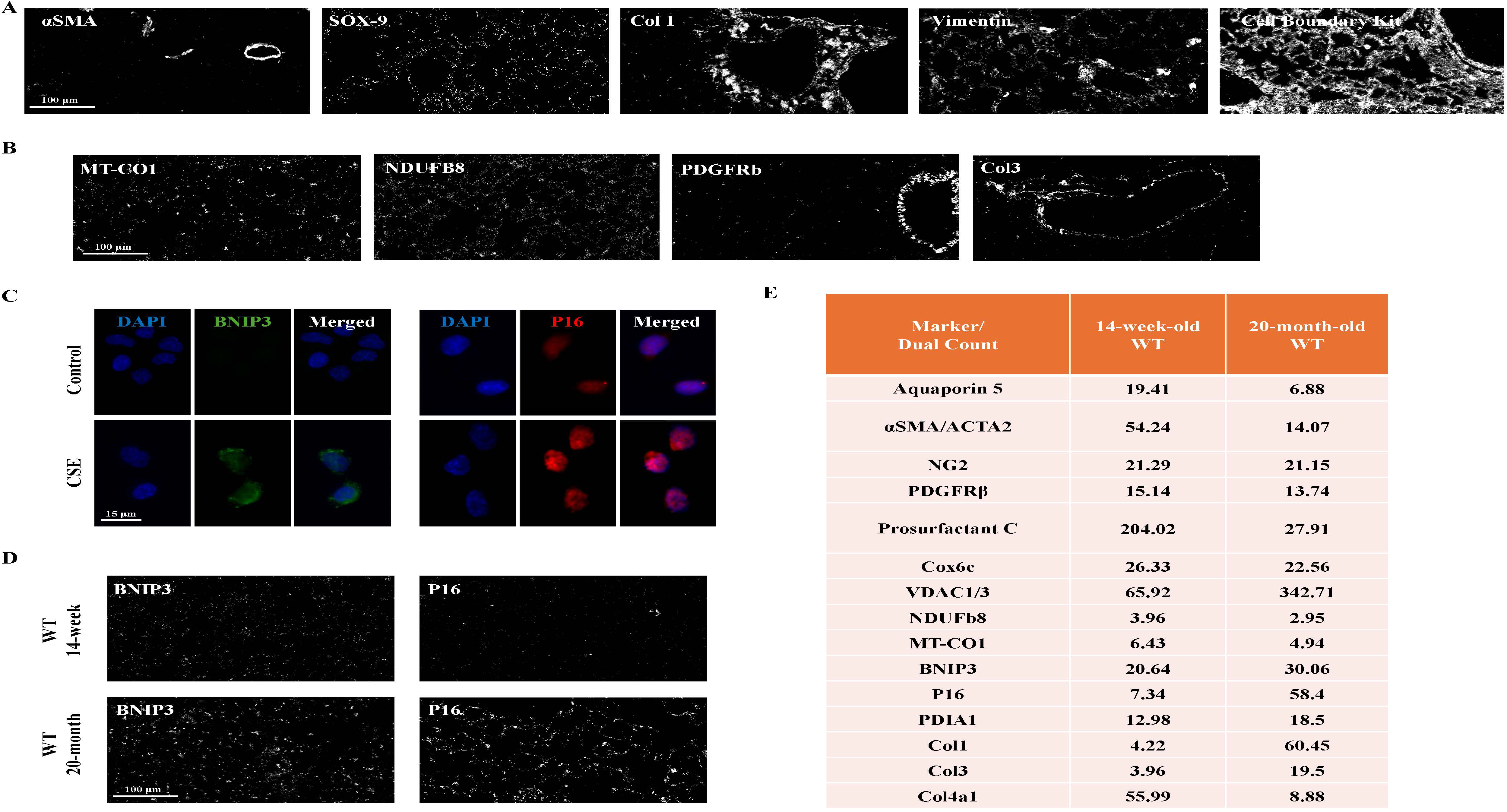
Verification of antibodies reactivity in lung tissues. **(A - B)** To validate antibody sensitivity and dynamic range, antibodies serial dilutions were tested in a biological condition expected to yield strongest signals. The highest dilution that gave a strong specific signal was chosen. Representative images of conditions used for further analyses are shown. (**A**) IMC images confirming lung reactivity of commercially available pre-conjugated antibodies against αSMA (^141^Pr), Vimentin (^143^Nd), SOX-9 (^147^Sm), Col1 (^169^Tm), and three cell membrane markers (^195^Pt/^196^Pt/^198^Pt), and (**B**) previously used for IMC of brain tissues but custom conjugated by us antibodies against MT-CO1 (^148^Nd), NDUFb8 (^174^Yb), PDGFRβ (^158^Gd) and Col3 (^152^Sm)**. (C)** Mouse Lung Epithelial 12 cell line was exposed (lower panel), or not (upper panel) to 6% CS for 7 hours to induce increase in BNIP3 (green) and p16 (red). BNIP3 and P16 were visualized using specified in Material & Methods antibodies. **(D)** 14-week and 20-month-old mice lung tissues were stained with anti-BNIP3 and P16 antibodies following their conjugation to ^173^Yb and ^175^Lu, respectively. ROIs were ablated using Hyperion IMC, and images were reconstructed in MCD viewer. Please note levels of BNIP3 and P16 were stress-or condition-specific. (**E**) Representative channel signal intensities (“Dual Count”) extracted using MCD Viewer. Data are representative of n=3 technical replicates (ROIs).

**Supplemental Figure 7.**
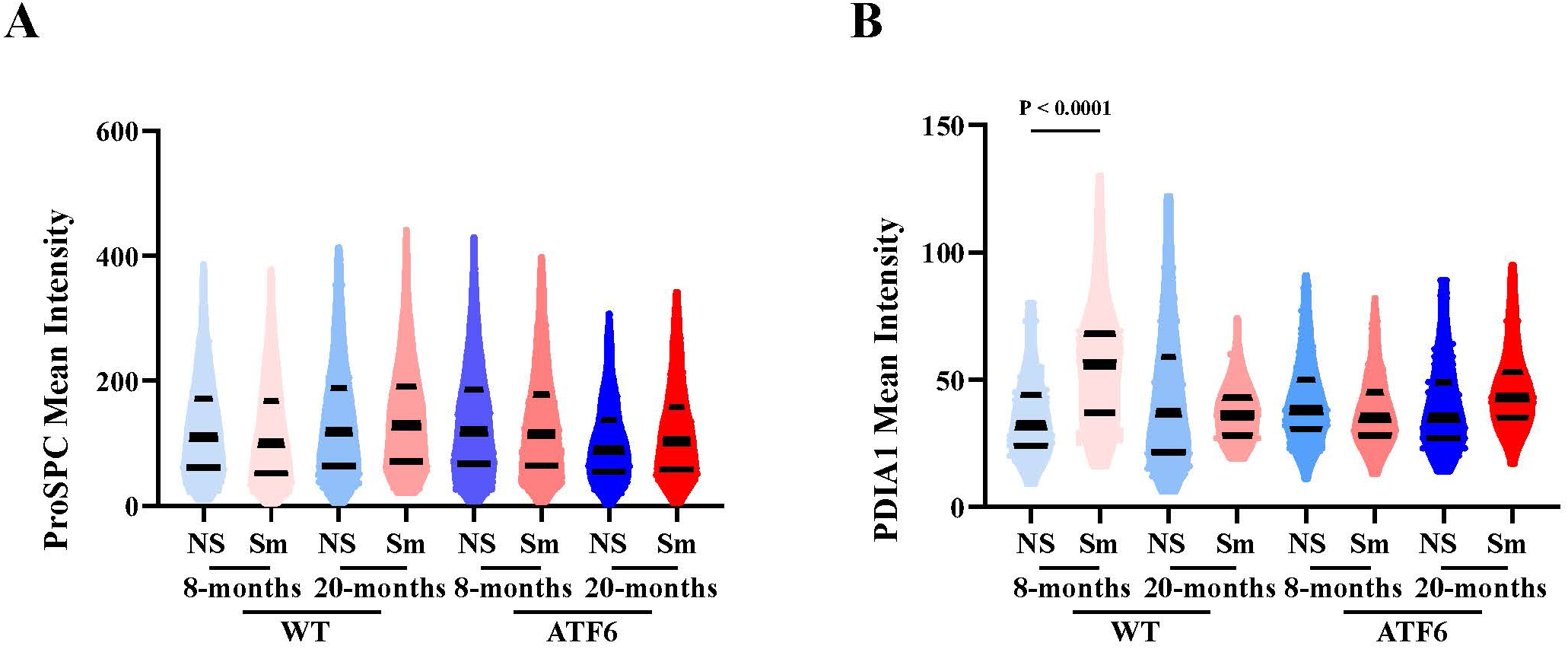
ATF6 deficiency does not alter surfactant or ER chaperone levels in AEC2-PopAs. Mouse lung parenchyma from 8-month-old and 20-month-old WT and ATF6-KO Sm or NS mice was profiled by IMC. AEC2 PopA were analyzed for protein levels of **(A)** pro-surfactant C; (**B**) PDIA1. Each violin plot represents analysis of >1000 segmented single AEC2s. Data are presented as mean ± SD and analyzed using one-way ANOVA with Tukey’s post hoc test; p < 0.05 was considered statistically significant.

**Supplemental Figure 8.**
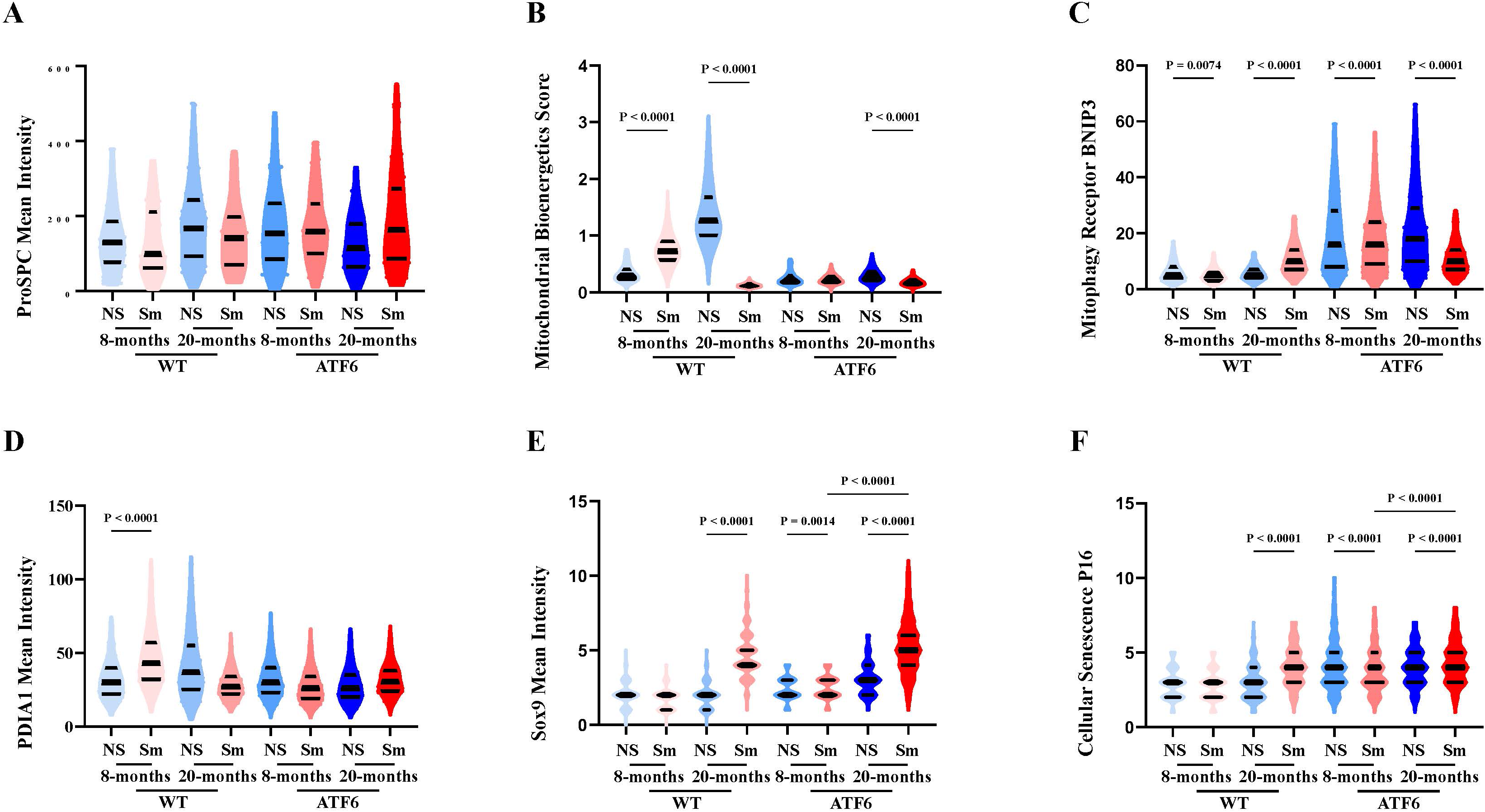
ATF6 deficiency abolishes adaptive increase in PDI-chaperone levels, and results in low mitochondrial bioenergetic score, increased mitophagy and senescence of AEC2-PopBs. Mouse lung parenchyma from 8-month-old and 20-month-old WT and ATF6-KO Sm or NS mice was profiled by IMC. AEC2-PopB subpopulation was analyzed for protein levels of **(A)** pro-surfactant C; (**B**) the mitochondrial bioenergetic score (Ndufb8, Cox6c, MT-CO1, and Vdac1/3) ; (**C**) mitophagy receptor BNIP3; (**D**) PDIA1; (**E**) Sox9; (**F**) p16. Each violin plot represents analysis of >1000 segmented single AEC2s. Data are presented as mean ± SD and analyzed using one-way ANOVA with Tukey’s post hoc test; p < 0.05 was considered statistically significant.

**Supplemental Figure 9.**
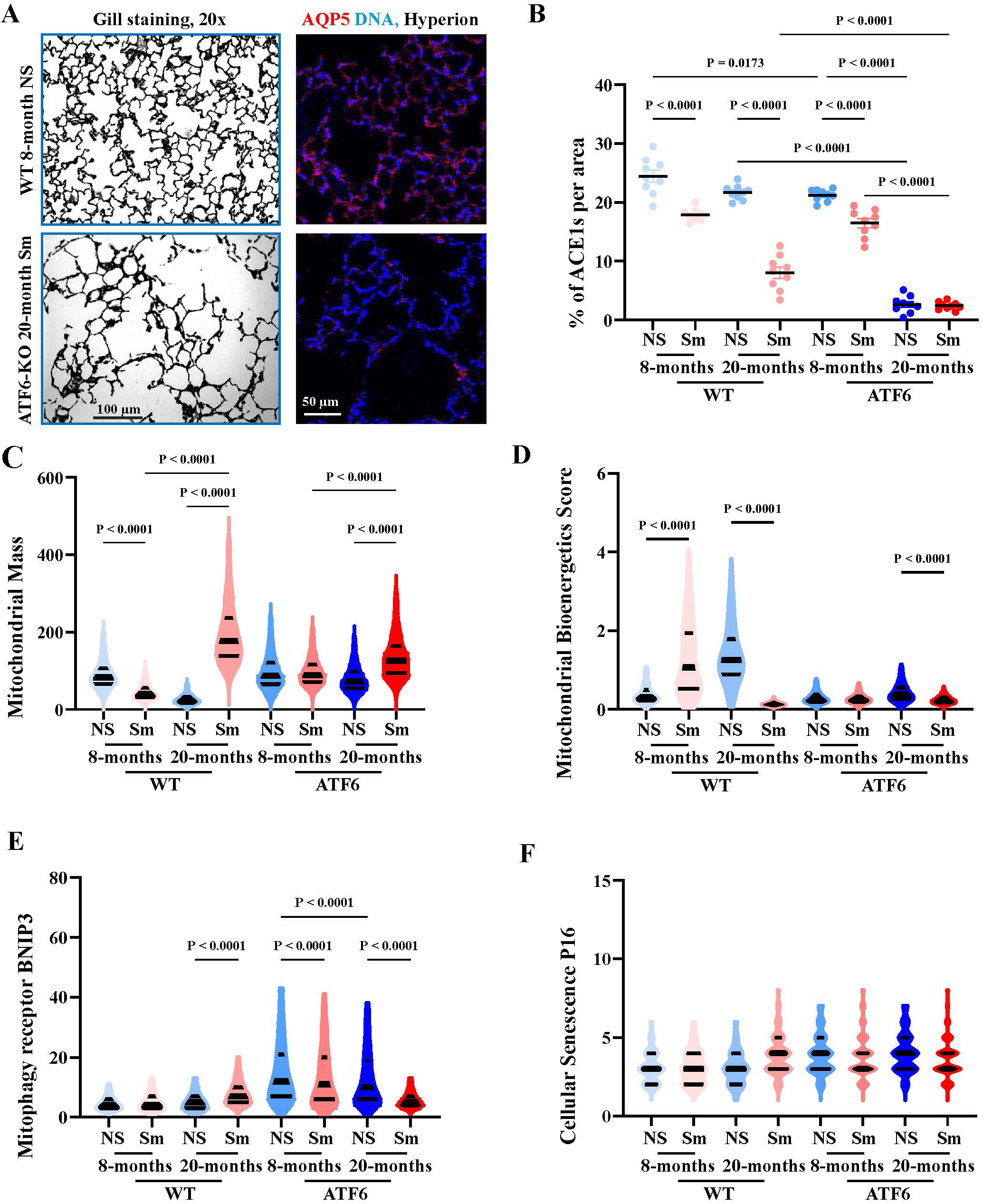
ATF6 deficiency impairs mitochondrial bioenergetics and increases mitophagy in AEC1s. Mouse lung parenchyma from 8-and 20-month-old WT and ATF6-KO Sm or NS mice is shown. **(A)** Gill’s staining (left). AQP5 (¹⁵³Eu, red) staining of respective to shown on the left conditions acquired by IMC, nuclei visualized by ⁹¹Ir/¹⁹³Ir and shown in blue (right). **(B)** Quantification of AEC1s abundance, expressed as the percentage of AEC1s per ROI, across experimental groups. Data are presented as mean ± SD from three biological replicates (n=3 mice per group), each with technical triplicates (3 ROIs per mouse), each dot represents one region analyzed. **(C -F)** Post-segmentation analysis of AEC1s for: **(C)** mitochondrial mass; (**D**) mitochondrial bioenergetic score; (**E**) mitophagy receptor BNIP3 levels; (**F**) cellular senescence marker P16 levels. Each violin plot represents analysis of >7500 segmented single AEC1s. Data are presented as mean ± SD and analyzed using one-way ANOVA with Tukey’s post hoc test; p < 0.05 was considered statistically significant.

**Supplemental Figure 10.**
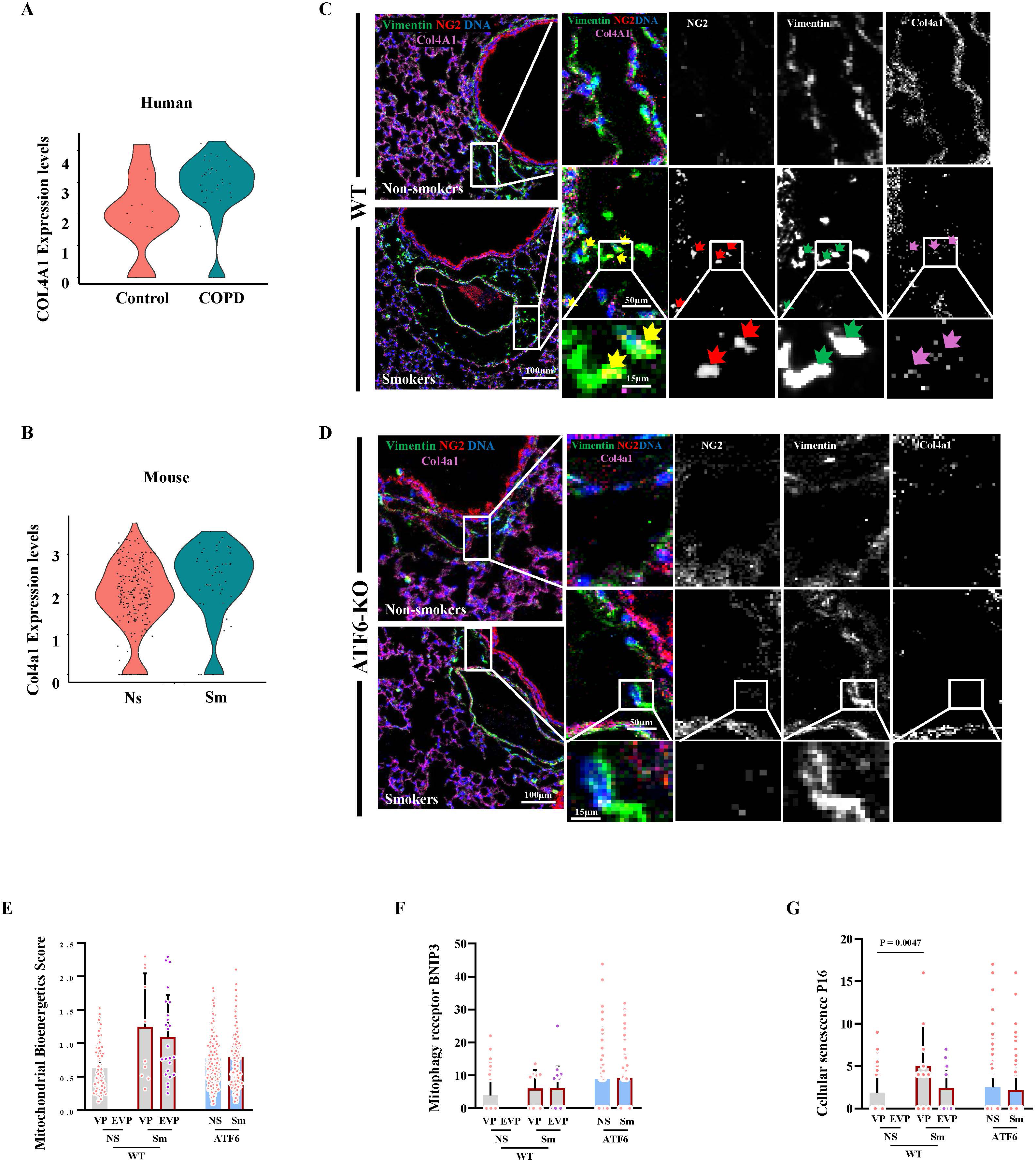
Characterization of cells within fibrotic lesions. IMC spatial analysis of lungs from WT and ATF6-KO mice exposed or not exposed to CS for 6 months. (**A - B**) Violin plots representing single-cell transcript levels of (**A**) COL4A1 in human pericytes and (**B**) Col4a1 in mouse pericytes. (**C - D**) NG2^+^ (^160^Gd) and vimentin^+^ (^143^Nd) cells in fibrotic lesions of WT (**C**) and ATF6-KO (**D**) mice. Enlarged areas from the left panels shown on the right. Boxed areas containing cells in the fibrotic lesion are further enlarged in the bottom panel. Arrows point to vimentin^+^NG2^+^ cells in the fibrotic areas. No vimentin⁺NG2⁺ cells were detected in the connective tissue around healthy airways and their associated blood vessels in WT healthy mice or in any ATF6-KO animals. Col4 was detected using ^154^Sm, and DNA was detected using ^191^Ir. NG2⁺PDGFRβ⁺ double-positive cells (pericytes) were classified into VP and EVP based on their location within or outside of pulmonary vasculature, as defined in Figure 6. Post-segmentation analysis of vascular and extravascular subpopulations of double-positive cells was performed. **(E)** Mitochondrial bioenergetic score was calculated based on the relative levels of ETC subunits and mitochondrial mass. **(F)** Levels of the mitophagy receptor BNIP3, and **(G)** levels of the cellular senescence marker P16. Each bar plot represents analysis of >800 segmented single pericytes. Data are presented as mean ± SD. Statistical analyses were performed using one-way ANOVA followed by Tukey’s post hoc multiple-comparison test; p < 0.05 was considered statistically significant.

## Material and Method

All raw data is provided in Supplementary Excel tables.

### Mice

C57BL/6 male and female mice (aged 8 - 10 weeks old) from Taconic Farms (Germantown, NY, USA) were used for experiments. ATF6-KO mouse was a gift from Dr. R. J. Kaufmann, where exon IV of ATF6α gene was deleted (1). All mice were housed and used for experiments under approved protocols of the University at Buffalo Institutional Animal Care and Use Committee (protocol PROTO202200042).

### Mice CS exposure in vivo

Mice were exposed to unfiltered cigarettes, twice a day, 5 days a week for 6 months (started from 8-10 weeks old or retired 14-month-old breeders) or to a sham, using the smoking chamber for the predominantly nose exposure (2, 3).

*Tissue processing and collection* was done as described (2, 4). Briefly, lungs were harvested 16 h post last CS exposure. Animals were terminally anesthetized by a combination of ketamine/xylazine (125/20 mg/kg) followed by cervical dislocation. Lungs were inflated by instilling intratracheally a 10 % neutral buffered formalin at 25 cmH2O for 10 min, ligated and removed. Inflated lungs were fixed in the 10 % formalin solution overnight, followed by 3 times, 15 mins of cold PBS wash and paraffin-embedded (**FFPE**). Serial sagittal sections of 4 μm were used for analysis. For all staining procedures, FFPE slides with lung tissue sections were baked at 60°C for 30 minutes, deparaffinized in xylene, and rehydrated through a graded ethanol series (100%, 95%, 70%, 50%, 30%, 1 min each) to distilled water. Antigen retrieval was performed using Target Retrieval Solution (Agilent Dako, S169984-2) for 30 mins at 96 °C.

### TUNEL staining

TUNEL staining was performed using the FragEL DNA Fragmentation Detection Kit (Cat #: QIA33-1EA, Sigma) according to the manufacturer’s instruction. Briefly, rehydrated section after antigen retrieval were incubated with the TUNEL reaction mixture, where **t**erminal **d**eoxynucleotidyl **t**ransferase (**TdT**) catalyzed the addition of labeled dUTPs to fragmented DNA strands, treated with a streptavidin-HRP conjugate followed by incubation with a DAB substrate to visualize apoptotic nuclei. Sections were counterstained with hematoxylin to provide structural context. Negative controls were maintained by omitting the TdT enzyme from the reaction mixture. Apoptotic activity was quantified by counting the total number of TUNEL^+^ cells across at least 10 fields per animal acquired at the magnification of 20x, using Leica DMi6 microscope brightfield. To account for variations in tissue density, results were normalized to ACL and expressed as the number of positive cells per mm of alveolar wall.

### Modified Gill lung tissue staining and Morphometric Analysis of Lung Airspace

Gill staining was performed as previously described in (5). Briefly, a custom staining solution was prepared by mixing equal volumes (125 mL each) of Gill’s stain (Sigma-Aldrich, St. Louis, MO; from Histology Core at SUNY Buffalo) and Harris Hematoxylin (Sigma-Aldrich; from Histology Core at SUNY Buffalo). Slides were incubated in the custom staining solution overnight. Following staining, slides were rinsed, differentiated in ammonium hydroxide solution (8 drops 6N NH₄OH in 250 mL diH₂O), dehydrated through a graded ethanol series (70%, 95%, 100%, 100%; 10 sec each), cleared in xylene for 1 min, 3 times, and mounted using Permount mounting medium (Fisher Scientific, cat # SP15-100). Stained slides were scanned at 20x magnification under Leica DMi6 microscope brightfield, 10 non-overlapping randomly chosen fields per slide were acquired by blinded to the slide identity personnel. Images were converted to grayscale (Black & White adjustment, -200% color influence) using Adobe Photoshop (Adobe Inc. (2024). Adobe Photoshop (25.0)) and saved in TIFF format without layers. ACLs were quantified using the deep learning-based tool Deep-Masker (https://mouse.pu-lab.com/home) (6). The default image scale (0.5) was utilized. To convert the output chord lengths from pixels to micrometers, a conversion factor was calculated based on the digital slide scale bar. A resolution of 0.667 µm/pixel was applied to the Deep-Masker output to generate final morphometric values in microns.

### Mac3 staining

Rehydrated sections after antigen retrieval were incubated with a primary antibody that recognizes MAC3 (Cat #: MCA2293GA, Bio-Rad), washed 3x with PBS, incubated with a biotinylated secondary antibody for 1 hour, washed 3x with PBS, incubated with streptavidin-HRP for 30 min, washed 3x with PBS. The chromogenic signal was developed using DAB (Cat #: ab64238, Abcam), counterstaining was performed with hematoxylin (cat #: H9627, Sigma). 10 non-overlapping serendipitously chosen fields per slide were acquired by blinded to the slide identity personnel using 20x magnification of Leica DMi6 microscope brightfield function. Digital images of Mac3-stained sections were analyzed using ImageJ (version 1.44, NIH). For each image, the quantification process was standardized using a custom macro to ensure reproducibility. Briefly, the procedure consisted of the following steps: 1. Color Deconvolution and Thresholding: Images were converted to the HSB (Hue, Saturation, Brightness) color space and split into individual channels. Segmentation of Mac3^+^ cells was performed by applying a threshold to the Brightness channel (range: 0–65) while maintaining full ranges for Hue and Saturation (0–255). A binary mask was generated by applying a logical AND operation across the HSB channels to isolate the low-brightness (darker) pigmented areas corresponding to the chromogen. 2. Morphological Filtering and Particle Counting: To identify individual macrophages and exclude non-specific staining or small artifacts, the Analyze Particles tool was employed. Objects were filtered based on a minimum size of 40 pixels and a circularity index of 0.53–1.00. 3. Data Output: The total number of Mac3-positive cells was automatically summarized for each field of view. To verify the accuracy of the segmentation, an overlay was generated by flattening the “Bare Outlines” of detected particles onto the original contrast-enhanced image. The mean number was determined and normalized to tissue density using ACL obtained by morphometry in formula: I = 0.3925/CL_(mean value given by Excel during morphometry)_ x10^3^ where, CL: Chord Length, I: amount of macrophages per mm lung tissue and 10^3^ adjustment to mm (7).

### Trichrome Staining

For the assessment of collagen deposition deparaffinized, rehydrated sections were incubated in Bouin’s fixative to intensify the final coloration. The sections were then stained with Weigert’s iron hematoxylin, followed by Biebrich scarlet-acid fuchsin and aniline blue treatments (Cat #: HT15-1KT, Sigma) to differentiate collagen fibers in blue from muscle and cytoplasm in red. To accurately evaluate perivascular and peribronchiolar fibrosis, the extent of collagen deposition was quantified by measuring the area of blue-stained fibers relative to the vessel perimeter in Fiji using “Freehand” function to highlight the area, then “measure” function was used to calculate the areas/perimeters. All 5 lobes of the section were scored, and all identified vessels that supplied terminal bronchi measured, 25-30 per animals, by investigators blinded to the identities of the experimental groups.

### RNA extraction and pathway-focused gene expression analysis

For gene expression analysis lungs were removed, flash-frozen, and stored at -80 ^◦^C for further analysis. Tissues, size of a couple of rice grains, were homogenized in 1ml of QIAzol reagent (Qiagen, Valencia, CA, USA). mRNA extraction from tissues was done using mRNeasy kits (Qiagen) as per manufacturer’s protocol.

*UPR-focused gene expression analysis* was performed using RT² Profiler PCR Arrays of Mouse UPR plates (Qiagen, 330231 PAMM-089A) according to manufacturer instruction. All Qiagen RT2 Profiler plates qRT-PCRs were performed using RT² SYBR Green ROX qPCR Mastermix (330520) on StepOne Plus Real-Time PCR system (Applied Biosystems). Results were processed by Qiagen algorithm software available at the data analysis web portal at http://www.qiagen.com/geneglobe. Samples were assigned to controls and test groups. CT values were normalized based on a Qiagen manual algorithm selection with Hsp90ab1 as a reference gene.

### scRNAseq data Acquisition and pre-processing

Publicly available scRNA-seq datasets were obtained from the GEO. Two distinct datasets were analyzed independently: *1. Human Dataset (GSE136831):* Comprising samples from COPD patients and controls. *2. Mouse Dataset (GSE168299):* Comprising samples from cigarette smoke-exposed mice and air-exposed controls. Raw gene expression matrices were imported into R and processed using the Seurat package. Quality control was performed to ensure data integrity.

### Cell Type Subsetting and Re-clustering

Cell type annotations and metadata available on GEO were utilized to identify major lineages. For this study, we specifically subsetted populations of AEC2 cells and pericytes. Following subsetting, the subpopulations underwent a secondary round of dimensionality reduction to refine the cellular structure: 1. **P**rincipal **C**omponent **A**nalysis (**PCA**): Performed on the variable features of the subsetted data (using RunPCA with npcs = 10). *2.* Neighbor Identification: A **S**hared **N**earest **N**eighbor (**SNN**) graph was constructed using the top 10 principal components (FindNeighbors). *3.* Dimensionality Reduction: **U**niform **M**anifold **A**pproximation and **P**rojection (**UMAP**) was calculated on dimensions 1:10 (RunUMAP) to visualize the specific substructure of these cell types.

### Differential Expression Analysis

Differential gene expression analysis was performed using the FindMarkers function in Seurat (default Wilcoxon Rank Sum test) to identify transcriptomic changes associated with disease or exposure. Comparisons were defined as follows: Human: COPD versus Control samples (Variable: Disease_Identity). Mouse: Cigarette smoke-exposed versus Non-smoker controls (Variable: smokers). DEGs were defined using the following criteria: *1.* Adjusted P-value: < 0.05 (Bonferroni correction). *2.* Expression Frequency: Genes were required to be expressed in a portion of cells in either group (default parameters were used). *3.* Fold Change: Average log2 fold change was calculated to determine upregulation or downregulation. Results were exported including p-values, average fold change, and percentage of cells expressing the gene in each group as CSV files.

### Functional enrichment analysis

To gain a deeper understanding of the biological processes associated with DEGs, a gene enrichment analysis was performed to examine biological processes and **K**yoto **E**ncyclopedia of **G**enes and **G**enomes (**KEGG**) pathways. Before analysis, a filter with p < 0.05 and fold change > 1.25 was applied to every DEGs for downstream analysis. The analysis was carried out using the g:GOSt function on the gProfiler web server (https://biit.cs.ut.ee/gprofiler/gost) (8). The significance threshold was set to the Benjamini-Hochberg **FDR** (**f**alse **d**iscovery **r**ate) value, and significant GO terms and KEGG pathways were defined by an adjusted p value of ≤ 0.05. For visualization purposes, bar plot representing the specific enriched GO terms and KEGG pathways were generated using the SRplot online server (https://www.bioinformatics.com.cn/en) (9).

### Ingenuity Pathway Analysis

DEGs from the human and mouse datasets were uploaded to the **I**ngenuity **P**athway **A**nalysis (**IPA**) software separately (10), using the expression log ratio and adjusted p values from the dataset as the observation. IPA’s Core Analysis function was used to gain a deeper understanding of altered signaling pathways in response to disease. The Canonical Pathways features from Core Analysis were used to see significantly affected pathways [absolute activation z score ≥ 2; −log_10_(p value) ≥ 2] on the basis of differentiation expression molecules.

### Targeted Network Construction

To investigate molecular connectivity, we defined specific gene groups of interest for network analysis. These groups included:

1. Human AEC2 DEGs associated with the GO term “cell differentiation” (GO:0030154).
2. Human AEC2 DEGs associated with inflammation cytokine pathways (identified via IPA Canonical Pathways).
3. Human AEC2_A DEGs associated with the programs that have been established in AEC2s to AEC1s transition.
4. Conserved Pericyte DEGs: Genes commonly regulated in both human pericyte and mouse pericyte datasets.
5. Conserved Pericyte DEGs associated with the GO term “extracellular matrix” (GO:0031012).
6. Conserved Pericyte DEGs associated with the GO term “cell differentiation” (GO:0030154).
7. Conserved Pericyte DEGs associated with collagen translation, trafficking, maturation, and deposition.
8. Conserved Pericyte DEGs associated with the programs that have been established in PTM.

Each group of common genes was input into IPA software separately. The “Path Explorer” tool was used to illustrate known relationships between genes. The “Grow” tool was used to expand the connections with a strict one-node limit to reveal the shortest molecular connections between the identified targets. Finally, the “Overlay” tool was utilized to map the expression data from the original datasets onto the generated networks, allowing for the visualization of gene expression patterns specific to each dataset (e.g., Human AEC2s or conserved Pericyte signatures).

### Tissue Staining for IMC

FFPE mouse lung tissue sections of 5 μm were used for IMC analysis. Sections were incubated at 60 ◦C for 2 h, deparaffinized in fresh xylene, and rehydrated through a graded ethanol series (100 %, 95 %, 80 %, 70 % v/v). Heat-induced epitope retrieval was carried out in citrate-based antigen unmasking solution (Vector Laboratories, 1X, pH 6.0) at 96 ^◦^C for 30 min using a steam chamber. To block non-specific binding, tissue sections were incubated in 3 % **b**ovine **s**erum **a**lbumin (**BSA**, sigma A7030) in PBS for 30 min at room temperature, followed by overnight incubation at 4 ^◦^C in a humidified chamber with 40 μL of the antibody cocktail diluted in 0.5 % BSA-PBS. The following day, slides were washed twice with 0.2 % Triton X-100-PBS and twice in PBS. Nuclei were counterstained with intercalator-Ir (see Table 1) for 30 min at room temperature, washed with double-distilled water, air-dried for at least 20 min and stored for IMC acquisition.

### IMC Panel Design and Antibody Labeling

Use of metal tags over fluorophores is preferable not only due to the increased number of proteins it allows to follow at once, but also since CS exposed tissues have high autofluorescence, which can limit their analysis using regular approaches. A twenty-markers IMC antibody panel was designed and validated to study proteins of interests and specific cell types of lung parenchyma distribution (**Table 1**). All custom antibodies were conjugated to metal isotopes using the Maxpar® X8 Multimetal Labeling Kit (Fluidigm) according to the manufacturer’s instructions.

### Antibody Verifying in lung tissues

20 metal conjugated marker panel was constructed (Table 1). Commercially available pre-conjugated antibodies against αSMA, Vimentin, SOX-9, Col1 and three cell membrane markers were purchased from Standard BioTools. Their performance in lung sections was confirmed/verified (**Sup. Fig. 6A**). Use of anti-PDIA1, anti-PrSPC, anti-AQP5, anti-Cox6c and anti VDAC1/3 antibody for lung IMC was previously published by us (11). Antibodies detecting MT-CO1, and NDUFb8 were previously used for IMC of brain tissues (12). We conjugated them de-novo and validated detection in lungs (**Sup. Fig. 6B**). Col3 and PDGFRβ were validated on lung slices after conjugation (**Sup. Fig. 6B**). BNIP3 and P16 were validated to give signals in MLE12 cells after 7 hours induced with 6% CS exposure (**Sup. Fig. 6C**) before conjugation. BNIP3 and p16 were validated on lung slices after conjugation (**Sup. Fig. 6D**). To ensure biological relevance, the entire panel was tested in lung tissues from 14-week-old WT mice and 20-month-old WT mice exposed to CS. Observed spatial and expression changes between these conditions aligned with established biological expectations, further verifying the sensitivity and accuracy of the conjugated antibodies. (**Sup. Fig. 6E**)

### IMC Data Acquisition

Imaging Mass Cytometry was performed using the Hyperion™ Imaging System coupled to a Helios™ mass cytometer (Fluidigm). Before laser ablation, panoramic optical images of each section were acquired at 20x magnification using the “Panorama” function of Hyperion system. For each animal, six regions of interest (ROIs; 500 × 500 μm each) were selected from the lung parenchyma. Three ROIs were chosen from emphysematous regions and three from fibrotic regions, guided by adjacent sections stained with Masson’s trichrome. Laser ablation was performed at a frequency of 200 Hz with a spatial resolution of ∼1 μm^2^/ pixel. Metal ion signals from antibody-labeled tissue were captured during ablation, preserving spatial context, and stored in .mcd format. Raw ion image data were visualized and exported using MCD Viewer (v1.0.560.6, Fluidigm).

### Cell Segmentation and Data Processing

Cell segmentation was performed using CellProfiler (version 4.2.6, 2021 Broad Institute). Composite images were first generated by merging DNA and membrane channels using the *ImageMath* module. A *GaussianFilter* was applied to smooth the membrane signal. Nuclei were segmented with the *IdentifyPrimaryObjects* module using global thresholding, a minimum–maximum diameter range of 4–12 pixels, and exclusion of objects touching image borders or falling outside the size range. Cytoplasmic boundaries were defined using the *IdentifySecondaryObjects* module with the “Watershed–Image” method, global thresholding (Minimum Cross-Entropy), a threshold correction factor of 1.0, and hole-filling enabled. Masks for both nuclear and wholecell regions were exported using *ConvertObjectsToImage* for downstream quantification.

### Marker Quantification Using Custom Software

A custom standalone application (IMC_Analysis) was developed to extract single-cell marker intensities from IMC images using single-cell segmentation masks. To enable automated, parallelized batch processing, raw single-channel TIFF images and their corresponding masks were organized hierarchically by main folder, experimental condition, murine subject, and ROI. Pixel quantification incorporated a customizable intensity thresholding mechanism to strictly isolate positive marker signals from background noise. By leveraging multi-core processing, the software evaluated each segmented cell to compute a comprehensive suite of metrics, comprising maximum intensity, mean and median positive intensities, positive pixel counts, and percent positivity, which were subsequently compiled into structured Excel workbooks. To facilitate specific downstream bioinformatic analyses, the mean positive intensity values were computationally extracted from these comprehensive results and exported as dedicated CSV files for each distinct ROI.

### Identification and Quantification of different cell types

Single-cell data extracted from IMC were imported into FCS Express 7 (version 7.24.0033; De Novo Software) in CSV format, as exported from custom software. For each ROI, a new layout was generated in FCS Express, and individual cell events were visualized using two-dimensional color dot plots. Marker expression levels were plotted using arcsinh transformed axes to enhance dynamic range and resolution of low intensity signals. Gating was performed manually using rectangular regions based on cell type specific marker expressions. A marker intensity threshold of 10^2^ (arcsinh-transformed) was applied to define cell-type specific marker-positive cells, based on alignment with corresponding spatial plots and confirmed visually against the original IMC images (MCD viewer), ensuring fidelity between spatial localization and marker expression.

Once gated, cell populations were color-coded and overlaid onto spatial scatter plots using “x-position” and “y-position” parameters to map the spatial distribution of different cell types within each ROI. Overlay features were used to confirm the accuracy of gating across different biological replicates and ROIs, maintaining consistent gate definitions. Gate statistics were generated to determine the proportion of different cells relative to the total cell population within each ROI. This percentage was used as a quantitative readout of cell type abundance. Results were consistent with previous findings and supported subsequent group-wise comparisons.

### Statistical Analysis for IMC

Following cell type classification, all subsequent analyses were restricted to cells expressing cell type specific markers of interest. Marker intensity values from gated cell type populations were exported from FCS Express 7 and analyzed using GraphPad Prism (version 10.4.1). Outliers were detected and removed using the **R**obust **R**egression and **O**utlier **R**emoval (**ROUT**) method with a false discovery rate (Q) of 1 %. One-way ANOVA was performed to assess differences across experimental groups. Variance homogeneity was evaluated using both Brown–Forsythe and Bartlett’s tests. Post hoc comparisons were conducted using Tukey’s multiple comparisons test. All statistical tests were two-sided, with adjusted P values < 0.05 considered significant.

### Mitochondrial bioenergetic score calculations

To assess mitochondrial bioenergetic functionality relatively to mitochondrial mass, a “Mitochondrial bioenergetic Score” was calculated for each identified individual cell. This score was defined as the average mean intensity of the OXPHOS subunits MT-CO1, NDUFb8, and Cox6c, normalized to mitochondrial mass, estimated by expression of the outer mitochondrial membrane porins VDAC1 & 3 using following formula - Mito Bioenergetic Score = (MT-CO1 + NDUFb8 + Cox6c) / VDAC1/3.

### Cluster analysis for IMC

Raw IMC intensity data were imported from Excel files into Python environment. 6 cell type specific markers, PrSPC, AQP5, PDGFRβ, NG2, αSMA and Col1 channels are included in the analysis. The raw marker intensity matrices from each condition were concatenated into a single dataset, and missing values were imputed with zeros. To stabilize variance and correct for the high dynamic range of IMC data, raw intensities were transformed using a log(x + 1) function. The log-transformed data were subsequently standardized using z-score normalization (scaling to zero mean and unit variance) via the StandardScaler function from the scikit-learn library. Single-cell analysis was conducted using the Scanpy toolkit (v1.11.4). Following feature scaling, we performed PCA to condense the data into five principal components. These top components capture the majority of the data’s variance while simplifying the feature space. A nearest-neighbor graph was constructed in PCA space using n_neighbors=30 and n_pcs=25. Non-linear dimensionality reduction was carried out using UMAP to visualize the cellular landscape in two dimensions. Unsupervised clustering was performed using the Louvain (13, 14) algorithm, applied at a resolution of 0.7. The analysis pipeline exports a comprehensive Excel file containing the annotated single-cell data (raw and log-transformed expression, UMAP coordinates, cluster assignments) and summary statistics, including mean marker expression intensities per cluster. Heatmap built in SRplot online servers was used to annotate each individual clusters.

## Notes

### Competing Interest Statement

The authors have declared no competing interest.

### Summary of Updates

Added funding information: "National Institute of Health/ National Heart Lung and Blood Institute (NIH)/(NHLBI) grant R01HL163168 to YB."

